# The regulatory landscape of optic fissure closure in the vertebrate eye

**DOI:** 10.64898/2026.02.11.705341

**Authors:** Brian Ho Ching Chan, Mariya Moosajee, Holly Hardy, James Prendergast, Joe Rainger

## Abstract

**Purpose:** Optic fissure closure (OFC) is a critical tissue fusion event in normal eye development and its failure results in ocular coloboma (OC), an inferonasal ocular tissue defect that persists throughout life. Most OC cases lack a genetic diagnosis, reflecting limited understanding of the genes and pathways that regulate OFC. We aimed to determine if OFC gene expression is regulated by the non-coding genome and identify novel candidate loci for OFC/OC.

**Methods:** Using the embryonic chicken eye, we performed integrated profiling of accessible chromatin and gene expression patterns during OFC. Matched RNA-seq and ATAC-seq from distinct spatial and temporal eye tissues were utilised with bioinformatic tools to identify regulatory genomic elements and infer gene regulatory networks, and to map synonymous loci in the human genome.

**Results:** We revealed domains of accessible chromatin during active fusion and the broader ventral retina that were distinct from those in the dorsal retina, implicating these loci for gene regulation during development and fusion of the optic fissure. *In silico* analysis using *de novo* motif enrichment and transcription factor (TF) foot-printing revealed TEAD, ZIC, and SOX TF activity during OFC, and retinoic acid signalling related TF activity in the dorsal eye. A subset of chicken OFC-specific peaks mapped to human cis-regulatory elements near known coloboma genes and OFC candidate genes identified in this study.

**Conclusions:** This provides the first evidence for dynamic cis-regulatory activity during OFC, identifies candidate loci for future mutational analysis, and offers new genetic leads for OC cases without a genetic diagnosis.

## BACKGROUND

Structural eye anomalies comprise a leading cause of childhood blindness, but despite a strong genetic disposition, most patients do not receive a genetic diagnosis^1–3^. Within these disorders, ocular coloboma (OC) is the most common and is considered as part of a clinical spectrum along with anophthalmia (no eye) and microphthalmia (small, underdeveloped eye), collectively referred to as MAC. Genetic causes of MAC manifest early in embryonic development^1,4^ and can occur in isolation or as part of a syndrome. OC is caused by a failure of epithelial fusion during optic fissure closure (OFC) in the ventral eye, leading to a persistent gap that can extend along the entire anterior-posterior length of the eye^5^, affecting the ciliary body, iris, neural retina, retinal pigmented epithelium (RPE), and choroid, and may extend to involve the macula and optic disc. It accounts for approximately 10% of childhood blindness and may affect up to 11.9 per 100,000 births^6,7^.

Next generation sequencing has improved MAC genetic diagnostic rates using whole genome sequencing and targeted gene panels, and approximately 33% of patients now have a genetic cause assigned^3^. In addition, zebrafish, chicken and mouse models of OFC have revealed candidate genes through transcriptional profiling of embryonic retina and optic fissure tissues that can be additionally included in genetic screening^8–10^. However, the diagnostic rates for MAC and in particular coloboma still remain low, prompting researchers and clinicians to consider causative genetic variants located in non-coding loci, poly-genic causes, or non-mendelian inheritance, and that MAC may be caused or influenced by environmental factors^4,11^.

Precedence for causative variants located in non-coding regions of the genome is emerging for structural eye malformations^12,13^, and have more recently been identified in patients with aniridia^14,15^, Axenfeld-Rieger Syndrome^16^ and MAC^17,18^. The low incidence of their discovery in cases of coloboma are likely to reflect the number and heterogeneity of genes associated with coloboma and because no attempts have so far been made to identify *cis* regulatory regions that are active during OFC.

We and others recently leveraged the experimental advantages of the chicken embryonic eye as a model for OFC^9,19–22^. The amount of actively fusing fissure makes it relatively straightforward to routinely and accurately extract stage-matched tissues from the fissure margins for profiling^9,20,22^. Genes expressed in the chicken optic fissure are similarly expressed in other species’ OFMs, including humans, and show conserved spatial and temporal expression patterns^9,22^. Chicken embryos have been used for mapping accessible chromatin across developmental stages and tissues to identify functional cis-regulatory elements (CREs), transcription factor binding sites (TFBS) and to determine gene regulatory networks (GRNs)^23^, and have provided novel data to reveal GRNs active in neural crest development^24^, paraxial mesoderm^25^, sex determination^26,27^ and gonadal development^28^. Thus, we hypothesised that mapping dynamically accessible chromatin in the chicken OFM during fusion-relevant stages could identify CREs and GRNs specific to OFC.

## METHODS

### Embryo incubation and dissection

Fertilised wildtype Hy-Line chicken eggs were incubated at 37°C. Embryos were developmentally staged according to Hamburger-Hamilton (HH) reference criteria^29,30^ and were collected at both stages HH30 and HH34, corresponding to the fusing and post-fusion stages of OFC respectively^20^. Fissure and dorsal tissues were manually dissected from each embryonic eye and were separately collected into cold Ringer’s solution (123.2 mM NaCl, 1.53 mM CaCl_2_, 4.96 mM KCl, 0.81 mM Na_2_HPO_4_, 0.15 mM KH_2_PO_4_ in H_2_O). Three biological replicates of each stage and type of tissues were prepared, where each sample contained pooled tissue for either dorsal or fissure eye collected from 10 embryos.

### ATAC-seq sample preparation and sequencing

Single cell dissociation of dissected tissues was conducted based on previously described methods^31^. 50,000 isolated cells were counted using a haemacytometer and were resuspended in PBS. Nuclei extraction, transposition reaction, and library preparation were all conducted following the previously described ‘omni-ATACseq’ method^32^. Briefly, the 50,000 cells were homogenised in cold ATAC lysis buffer (10 mM Tris-HCl pH 7.5, 10 mM NaCl, 3 mM MgCl_2_, 0.1% NP-40, 0.1% Tween-20 and 0.01% digitonin). Nuclei were isolated by centrifugation at 500 RCF for 10 minutes at 4°C. Transposition was conducted using Tn5 transposase and TD buffer (Illumina, Cat no 20034197). Transposed DNA was cleaned using a MinElute Reaction Cleanup Kit (Qiagen, Cat no 28204), and then partially amplified by PCR using NEBNext High-Fidelity 2X PCR Master Mix (NEB, Cat no M0541) with i7 reverse indexing primers each with a unique barcode together with generic i5 forward primers were used to label each sample. Incubations at 72°C for 5 minutes and 98°C for 30 seconds, followed by five cycles of 98°C for 10 seconds, 63°C for 30 seconds and 72°C for 1 minute were used for amplification. The number of additional amplification cycles was determined by qPCR using Luna Universal qPCR master mix (NEB, Cat no M3003), where the additional cycles was determined as the number of cycles required to reach ¼ of the maximum intensity in qPCR. Libraries were cleaned using AMPure XP beads (Beckman Coulter, Cat no A63880). Library concentration was quantified using Qubit dsDNA High Sensitivity Assay Kits (Invitrogen, Q32851), while library quality was determined using 4200 Tapestation System (Agilent). 75bp paired-end sequencing of pooled libraries was then conducted using the Nextseq 2000 platform (Illumina), with at least 180 million reads generated per sample (**Supplemental Table S1**).

### ATAC-seq Alignment and Peak Calling

Adaptor sequences were detected and cut out using Trimgalore! and Cutadapt v1.16^33^. Sequence quality pre- and post-trimming was evaluated using FastQC (https://www.bioinformatics.babraham.ac.uk/projects/fastqc/) version 0.11.4. Read pairs were aligned to the Ensembl chicken reference genome (galGal6) to generate binary aligned map (BAM) files using Bowtie2 version 2.3.4.3^34^ setting the maximum inferred fragment length to 1.5kb with the following parameters: --very-sensitive -X 1500 -k 10. Quality control metrics for the individual sample, including the percentage of duplicated and mitochondrial reads, were generated using ataqv version 1.2.1^35^. To call regions of significantly open chromatin, Bowtie aligned BAM files of technical replicates were merged, sorted by query name and indexed using Samtools version 1.6^36^. Subsequently, peaks of each condition were called from the corresponding biological replicates using the Genrich pipeline version 0.6^37^ with the following parameters: -j -r -e MT -q 0.05. BAM-converted bigWig tracks were visualised using pyGenomeTracks version 3.6^38^.

### Differential peak analysis

Duplicated and mitochondrial reads from BAM files were removed using Picard version 2.26.0 and Samtools version 1.16^36^ respectively. All peak coordinates called by Genrich across different conditions were merged using BEDtools version 2.30.0^39^. Non-duplicated read fragments of each sample within the merged list of peak coordinates were counted by featureCounts in paired end mode (-B -p) to generate a count matrix^40^. To identify differentially accessible (DA) peaks, pairwise comparison was conducted using DESeq2 version 1.42.0^41^. Genomic loci with a Benjamini–Hochberg adjusted P-value< 0.05, and a log_2_ fold change > 0.5 or < -0.5 were marked as DA peaks in each comparison. Gene annotations of DA peaks was based on Ensembl *Gallus gallus* (version 105), and added using GenomicsFeatures version 1.54.3^42^ and ChIPSeeker version 1.46.1^43^.

### Motif analysis and transcription factor occupancy analysis

*De novo* motif enrichment analysis was performed using HOMER version 4.11^44^. The findMotifsGenome.pl function from the HOMER package was used to identify *de novo* motifs over-represented in the given list of peaks with the following parameters: -size 200 -mask. To determine motif density of TF in a given set of peaks, annotatePeaks.pl from the HOMER package was used with the following setting: -size 1000 -hist 5, which refers to searching against a total window size of 1000bp (i.e. +/- 500bp in 5’- and 3’-orientations) and a bin size of 5bp. To identify the transcription factor occupancy, ATAC-footprinting of the ATAC-seq dataset was conducted using TOBIAS^45^. In short, the TOBIAS standard workflow was divided into three steps: (i) Tn5 bias correction from ATAC data, (ii) calculation of footprint score at peak regions, and (iii) prediction of differential binding of transcription factors in two different sets of peaks^45^. The numbers of motifs supplied to TOBIAS were 5,272 cis-BP 2.0 inferred chicken motifs and 1,912 JASPAR 2024 vertebrate motifs^46,47^. Aggregate plot values were plotted using the PlotAggregate function of TOBIAS with the following parameters: --share-y both --plot-boundaries.

### CTCF dataset processing

Publicly available chicken CTCF-ChIP data^48^ were downloaded using SRA NCBI toolkit. Adaptor trimming was done using Trimgalore! and Cutadapt v1.16^33^. Alignment to the galGal6 genome using Bowtie2 was done using Bowtie2 version 2.3.4.3^34^. Peak calling with a q-value threshold of 0.05, and normalisation to input control was done using Genrich version 0.6^37^ with the following setting: -c -y -r -e MT -v - q 0.05.

### Determining proximal peaks of gene of interest

To obtain a list of differentially expressed genes of interest in the different comparisons, a count matrix of stage-matched RNAseq data was obtained from Trejo-Reveles *et al.* (2023)^9^ using the SRA NCBI toolkit. Differential expression analysis was conducted using DESeq2 version 1.42.0^41^. Peaks located within 20kb upstream of a transcription start site (TSS) and 20kb downstream of the end position of the gene of interest were marked as proximal peaks. For distal peaks, a topologically associated domain (TAD) boundary was first predicted using constitutive CTCF sites, which marked putative insulator positions^49^, based on the publicly available chicken CTCF-ChIP dataset. The TAD boundary of the given gene was defined by the highest CTCF bound score, determined by Genrich, within 250kb upstream and downstream, respectively, of the gene. If no CTCF bound site was found within 250kb upstream or downstream of the gene, the search window for a CTCF bound site would be extended to 500kb accordingly.

### RNAseq-ATACseq datasets correlation

TPM values from stage matched chicken embryonic eye RNA-seq data from Trejo-Reveles *et al.* (2023)^9^ were obtained using the SRA NCBI toolkit. The dataset was processed and analysed as described in Trejo-Reveles *et al.* (2023)^9^. To determine transcriptional start site (TSS) accessibility, non-duplicated DNA fragments located in a window 1000bp upstream and downstream of all non-mitochondrial protein-coding genes recorded in Ensembl (version 105) were counted using featureCounts^40^. Paired end reads with both ends mapped, and with overlapping reads not discarded, were counted using featureCounts.

### Identification of conserved enhancers in the human genome

Genomic sequences of putative OFC enhancers were extracted from the chicken galGal6 FASTA file. These DNA sequences were lifted over (aligned) to the human GRCh38 genome using NCBI BLAST+^50^ via a custom Python script with an E-value threshold of < 1 x 10^-5^. The identified human genomic regions were queried against the ENCODE Search Candidate CREs (SCREEN; Registry V4) to determine whether enhancer activity for these loci had been previously annotated in humans^51^. Only CREs labelled ”proximal enhancer”, ”distal enhancer” or ”promoter” by SCREEN were included in the analysis. A one-sided Fisher’s exact test was used to determine if the ENCODE-annotated human CREs were enriched within the lift-over peak set. The background DNA sequences for the statistical test were 4,029 randomly selected chicken genomic regions that were matched to the test peaks of interest (i.e. chicken OFM sequences) in terms of GC content and length, and mapped to the human genome using NCBI BLAST+ alignment. GC content was determined using nucBED from the Bedtools suite^38^.

### Immunostaining and *in-situ* hybridisation

Embryos were fixed with 4% paraformaldehyde (PFA) for 4 hours at room temperature, followed by 10% sucrose in PBS at 4°C overnight then frozen-embedded in Neg-50 (Epredia) and stored for up to 6 months at -80°C. Cryosectioning was performed with a Leica CM1900 Cryostat with sections cut at 14μm thickness. *In-situ* hybridisaiton using DIG labelled RNA riboprobes was conducted as described previously with the following gene specific primers for PCR-based amplification and synthesis of *TEAD1* riboprobes: Fwd 5’-AGAACAGGGAAGACACGGACC – 3’ and Rev 5’ -GTGTCCACGGATTCAAGCAACG -3’ and T7/T3 sequences added for *in vitro* transcription^20,52^. Immunofluorescence of YAP1 was performed using methods described previously^20^, but using rabbit anti-human YAP1 antibody (D8H1X, Cell Signalling Technology) diluted 1:50 in block solution (PBS, 0.1% BSA, 0.05% Triton X-100) and incubated overnight at 4°C, and detected with Alexa-fluor 488nm goat anti-rabbit secondary antibody (1:1000 dilution; A11037, Thermo Fisher) and DAPI (1:1000 dilution; D3571, Thermo Fisher).

## RESULTS

### ATAC-seq in the developing chicken eye

To identify dynamic chromatin associated with the progression of fusion at the optic fissure, we prepared dissected chicken ocular tissue samples for ATAC-seq (**Figure 1A**). In the chicken embryo, optic fissure closure begins around embryonic day 6-7 (HH29-31) and continues for approximately 60+ hours^20^ but is fully fused by HH33-34. Therefore, a HH30 OFM contains open, fusing and fused regions of retina whereas a HH34 OFM is completely fused (**Figure 1B**). At these stages, the dorsal retinal tissue is highly similar to the ventral retina in its broad anatomy but has no fusion event, so is a useful contrast against the OFM to identify fusion-associated gene expression dynamics^9,20,53–55^. We therefore dissected OFM and dorsal retina tissues at these two stages in 3 replicates (n=12 total samples) for ATAC-seq for comparative analyses to reveal chromatin dynamics during OFC (**Figure 1C**). Nucleosomal profiling of reads indicated clear spikes at approximately 50 bp, 180-240 bp, and 315-470 bp, corresponding to the periodicity expected from free DNA, mono-nucleosomal, and di-nucleosomal DNA fragments, respectively^56^ (**Figure S1A**). Reads were then mapped to galGal6 and uniquely mapping reads (73%) versus duplicate reads (20%) and unmapped reads (2%) were determined (**Figure S1B**). Of the mapped reads, 1% were mapped to the mitochondrial genome and 65% mapped to autosomes (**Figure S1C**). Unmapped, duplicate and mitochondrial reads were removed from further analysis. Sample-to-sample variance was determined using Euclidean distance and principal component analysis (PCA) (**Figure 1D**, **Figure S1D-E**), which confirmed that samples clustered according to biological conditions: temporal identity (HH30 or HH34) for PC1 (67% variance), and spatial identities (fissure or dorsal) for PC2 (16% variance). PC1 and PC2 together accounted for 83% of the total variance in samples (**Figure 1D**, **Figure S1D**). Similarly, Euclidean distance analysis confirmed that inter-sample variation reflected underlying biological differences (**Figure S1E**).

**Figure 1.**
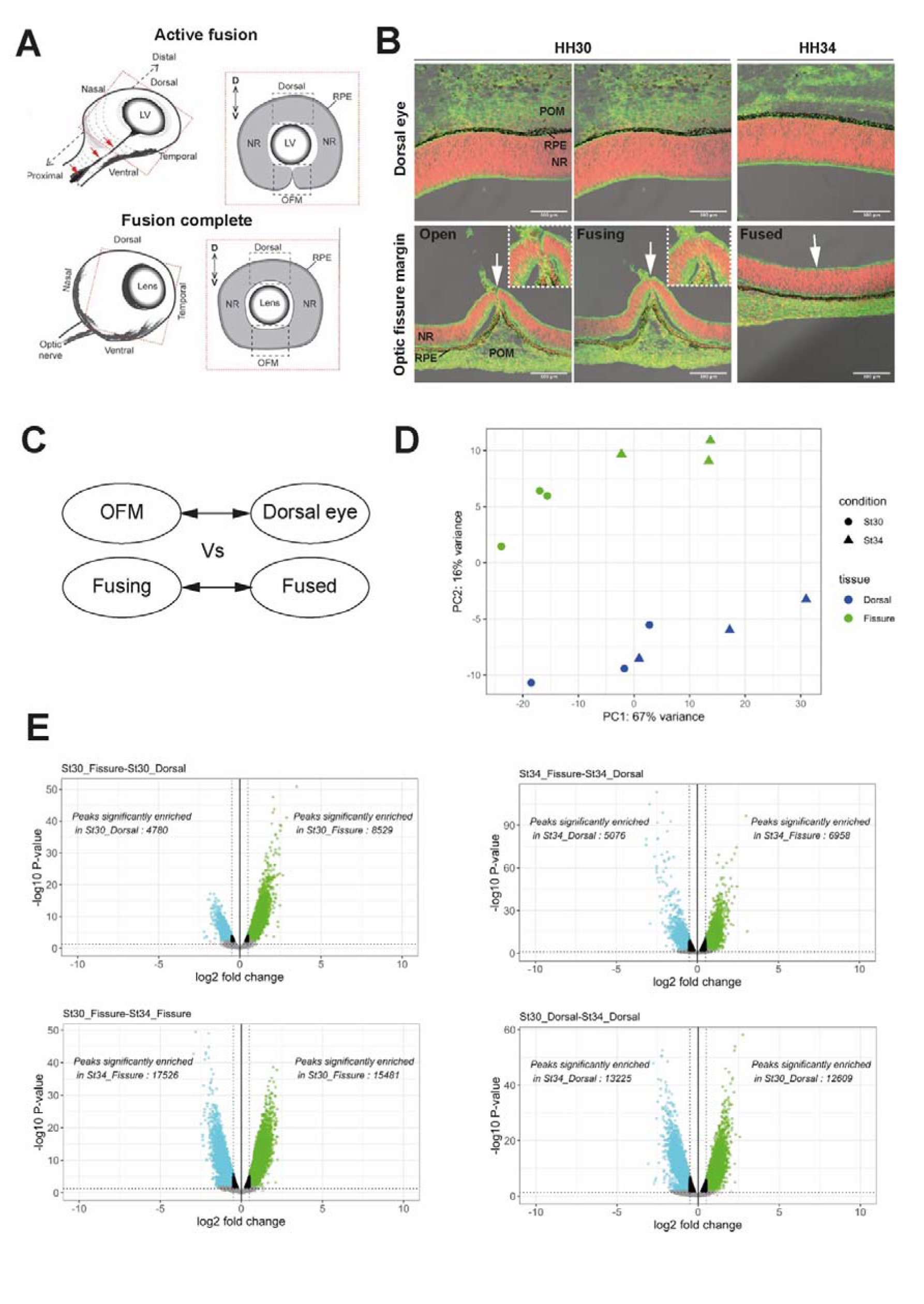
Profiling of genome-wide open chromatin during optic fissure closure. (**A**) Schema representing the optic fissure (arrows) in the ventral retina during vertebrate eye development, with virtual sagittal sections to illustrate the fusion of the fissure (boxes). Hatched “dorsal” and “OFM” boxes indicate regions dissected for ATAC-seq. (**B**) Sagittal cryosections of the chicken embryo illustrating active fusion (HH30/embryonic day 7) and fully fused (HH34/E9). Sections are stained with phalloidin (green) for F-actin, DAPI (red) for cell nuclei, and brightfield images to indicate the retinal pigmented epithelium (RPE, black). Arrows indicate fusion point in open and fusing (HH30) OFM, and equivalent region in the fused OFM (HH34). (**C**) Comparison schema for differential chromatin accessibility analysis: active fusion (HH30) optic fissure margin (OFM) versus fused (HH34) OFM, alongside inter stage spatial comparisons (OFM versus dorsal dissected tissue). (**D**) Principal component analysis (PCA1 and PCA2) of mapped reads for each sample group (HH30 and HH34, dorsal and OFM; 4x groups in total, 3 replicates per group). (**E**) Volcano plots showing DA peaks for each pairwise comparison. Log 2-fold change and log 10 P-value of each peak coordinate was determined using DESeq2. The number of significant DA peaks identified in each condition is annotated. Vertical dotted line represents the log 2-fold change cut offs of 0.5 and -0.5; horizontal line indicates adj-*P* 0.05. Abbreviations: NR, neural retina; POM, periocular mesenchyme. [Panel **A** has been amended from Chan et al., 2021 ^5^].

### Identification of spatial and temporal differentially accessible chromatin

We then proceeded to determine differentially accessible (DA) chromatin regions to map candidate regulatory elements for genes important for OFC. Comparisons were: HH30 fissure vs HH30 dorsal, HH30 fissure vs HH34 fissure, HH34 fissure vs HH34 dorsal, and HH30 dorsal vs HH34 dorsal (**Figure 1C**). First, all peak coordinates identified across the different conditions were merged to obtain a single list of coordinates for subsequent evaluation of differential chromatin accessibility. The distribution of locations within gene architecture for these peak coordinates across the different conditions were highly similar (**Figure S1F**). Most peaks were in distal intergenic regions (∼36% for all sample groups) or introns (∼40%), consistent with where enhancer regions are typically located. A further 16-17% were located tightly at transcriptional start sites (TSS) regions (**Figure S1F-G**), consistent with achieving high signal to noise ratios for our samples ^56,57^. Reads for each peak were then used to generate a count matrix, and paired combinations were tested for differential accessibility by pairwise comparisons using DESeq2 with adjusted P-value (adj*P*) < 0.05 and absolute log_2_-fold change (L2FC) >0.5. We found the greatest differences were observed temporally between equivalent tissue regions at different developmental stages (**Figure 1E**, **Table 1**). In contrast, the spatial comparisons had fewer total DA peaks, but it was noticeable that the HH30 OFM had markedly more DA peaks than the dorsal DA peaks.

**Table 1.** Number of ATAC-seq peaks with differential accessibility for each comparison.

| Pairwise comparison (# significant peaks) | Total number of DA peaks |
| --- | --- |
| HH30 OFM (8,529) vs Dorsal (4,780) | 13,329 |
| HH34 OFM (6,958) vs Dorsal (5,076) | 12,034 |
| HH30 OFM (15,481) vs HH34 OFM (17,526) | 33,007 |
| HH30 Dorsal (13,225) vs HH34 Dorsal (12,609) | 25,834 |

### Correlation analysis of matched ATAC-seq and RNA-seq

We then sought to assign these DA peaks to genes involved in OFC. To do this we integrated our pre-existing RNA-seq data collected from equivalent stage tissues in the chicken eye (HH30 & HH34; dorsal and OFM)^9^. First, we performed comparable differential gene expression (DEG) analysis to match our ATAC-seq analysis for the conditions (**Figure 1C**). In combination, all four pairwise comparisons revealed genes with significant differential expression (adjP < 0.05, L2FC >1 or <-1) (**Supplemental Figure S2A; Supplemental Table S2**), and we observed greater numbers of DEGs across the temporal (HH30 vs HH34) comparisons than we did for the spatial comparisons (fissure vs dorsal), which reflected our ATAC-seq data.

We then examined if the RNA-seq and ATAC-seq datasets correlated such that a combined analysis from the two datasets would be valid. Correlation was performed by matching ATAC-seq TSS accessibility to stage-matched RNA-seq transcript counts for each known coding gene. The Spearman correlation between TSS read counts and the RNA-seq counts for the corresponding coding genes ranged between 0.56 to 0.6 (*P* <2.2×10^-26^) at all four biological conditions examined, suggesting a moderate overall correlation between chromatin accessibility and gene expression^58^ (**Figure 2A**; **Supplemental Figure S2B**) and consistent with previous similar studies^59,60^. The density plots suggested a bi-phasic pattern of correlation, with two separate events that related to high and low expression levels: most genes with low transcript expression levels (log_10_(TPM+1) <1) had varying levels of promoter accessibility whereas for higher expressed genes TSS accessibility was consistently higher. This indicated a small degree of correlation among lowly expressed genes in the datasets, but a higher correlation for OFM enriched genes during fusion.

**Figure 2.**
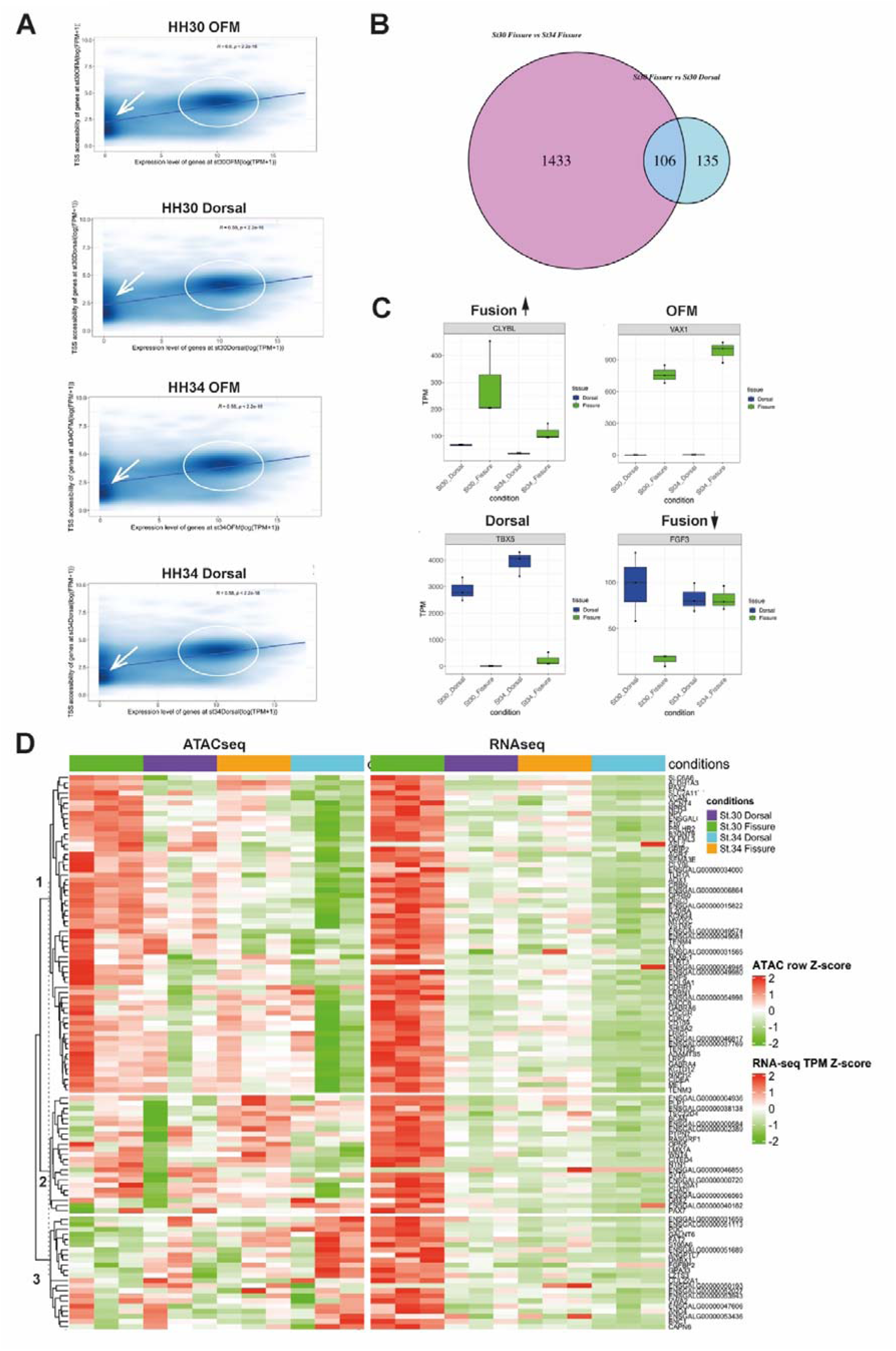
Correlation of ATAC-seq and RNA-seq datasets. (**A**) Density plots showing the distribution of expression levels and corresponding promoter accessibility of each coding gene at HH.30 fissure, HH.30 dorsal, HH.34 fissure, and HH.34 dorsal. Gene expression level for each gene is shown as log10(1+Transcript per million), while the open chromatin accessibility level is shown as log10(1+Fragments per million). R-value from Spearman correlation and the corresponding P-value are shown. Arrows indicate correlation for low expressed genes [log10(1+Fragments per million) approx. £1], and ellipses indicate correlation for genes with higher expression levels [log10(1+Fragments per million) approx. ³10]. (**B**) Venn diagram indicating fusion specific DEGs determined by comparison between HH30 OFM versus HH30 dorsal and HH34 OFM DEGS. (**C**) Boxplots of gene expression values (TPM) for representative genes displaying context specific differential expression. (**D**) Combined ATAC-seq and RNA-seq Z-score heatmaps showing k-means clustering outputs for correlation of TSS accessibility and gene expression for 106 fusion enriched DEGs, with dendrogram and cluster labels shown on left.

We then performed multiple comparisons to determine DEGs that were: (i) specifically enriched during fusion (“fusion specific” OFM > Dorsal and HH30 > HH34), (ii) repressed during fusion (Dorsal HH30 and HH34 and HH34 OFM > HH30 OFM), (iii) “OFM enriched” at both stages (OFM > Dorsal), or (iv) were dorsal enriched eye at both stages (Dorsal > OFM), and this analysis identified known regionally and temporally regulated genes, including 106 fusion specific DEGs (**Table 2**; **Figure 2B**; **Supplemental Figure S2C**). Example gene expression profiles for these DEG categories are shown in **Figure 2C** and included *CLYBL* as a fusion specific gene whose expression was previously reported in OFM pioneer cells prior to and during fusion but was then rapidly downregulated in the fused OFM^52^. Conversely, *FGF3* is a fusion repressed gene, and previous studies have shown that its expression is specific to cells in the central retina and is important for retinal ganglion cell differentiation^61^. *TBX5* was within the dorsal enriched DEGs category, and its expression is specific to chicken embryonic dorsal retinas and absent from ventral regions^62^. *VAX1* was a OFM enriched gene and previous studies in a range of vertebrates have shown it to be persistently expressed in the ventral retina throughout optic cup development and after OFC has completed^9,63^.

**Table 2.** Number of DEGs specific to each category.

| Category | Number of significant DEGs |
| --- | --- |
| Fusion enriched | 106 |
| Fusion repressed | 164 |
| OFM enriched | 87 |
| Dorsal enriched | 49 |

### Correlation of promoter accessibility to fusion enriched DEGs

Given our interest in understanding the regulation of fusion specific genes, we next determined whether promoter accessibility patterns matched transcript expression patterns in the 106 fusion specific DEGs. *K*-means clustering separated these into 3 clusters according to their TSS accessibility (**Figure 2D**): 62 of the 106 genes (57.9%) belonged to cluster 1, in which their TSS showed fusion specific accessibility that correlated to their expression profile. Specifically, these genes had the highest TSS accessibility for HH.30 OFM. Cluster 2 comprised of 22 genes (21.5% of all fusion specific enriched genes) with partially matching accessibility, in which accessibility was generally high within OFM HH.30, OFM HH.34 and dorsal HH.30 but low for dorsal HH.34. Cluster 3 comprised 21 fusion specific genes (19.8% of all fusion specific enriched genes) with non-matching TSS accessibility that was highest for dorsal HH.34 and generally low for OFM HH.30. These results indicate that for our data, open chromatin regions around TSS generally reflected the gene expression levels and therefore is likely to enable detection of regulatory elements for fusion specific DEGs.

### Identification of fusion specific loci with differentially accessible chromatin

We next attempted to associate DA peaks with the 106 fusion specific DEGs to identify regulatory networks in OFC. Enhancers may exist at various ranges and can be >400kb from the TSS of the coding genes they regulate^24,64–66^, however, most are not functionally characterised or easily mapped to target genes. To begin with, we used conservative search parameters of including all introns and exons (40kb total mean gene length) together with sequences 20kb upstream of the TSS and 20kb downstream from the 3’-UTR for each of the fusion specific DEGs, which identified a total of 1504 peaks for all conditions (**Supplemental Figure S3A**; **Supplemental Table S3**). Most gene-associated peaks were situated in promoters, distal intergenic, or intronic regions (**Supplemental Figure S3C**), with a median proximal peak length 426bp. Of these, 803 displayed significant differential accessibility in at least one of the pairwise comparisons, with 105 of these DA peaks classified as fusion specific and 139 peaks that were OFM enriched (**Table 3**). To expand our search for DAs, search windows were increased to +100kb flanking each fusion specific DEG (gene length +200kb) or were determined by topologically associating domain (TAD) boundaries for each DEG using publicly available chicken embryo CTCF-ChIP data^48,67^. An average TAD boundary size of 288kb was observed across all fusion specific genes (**Supplemental Figure S3B**), consistent with previous CTCF loop distance averages^68,69^. Overall, this increased the discovery of condition associated peaks, with 234 fusion specific DA peaks and 295 OFM enriched peaks identified (**Table 3**). The 10 highest fusion specific DA peaks are listed in **Table 4**. In conclusion, our matched RNA-seq and ATAC-seq analyses identified plausible enhancer regions with differential accessibility that correlated with spatial and temporal differential gene expression during OFC and was able to specifically identify DA loci associated with fusion specific gene expression.

**Table 3:** Peaks identified associated with fusion-specific DEGs in each condition of enrichment using size-based or TAD determined search windows.

| Interval size | OFM enriched | Dorsal enriched | Fusion enriched | Fusion repressed |
| --- | --- | --- | --- | --- |
| 40 kb | 139 | 9 | 105 | 10 |
| 200 kb | 227 | 32 | 188 | 47 |
| TAD boundary | 295 | 30 | 234 | 41 |

**Table 4.**
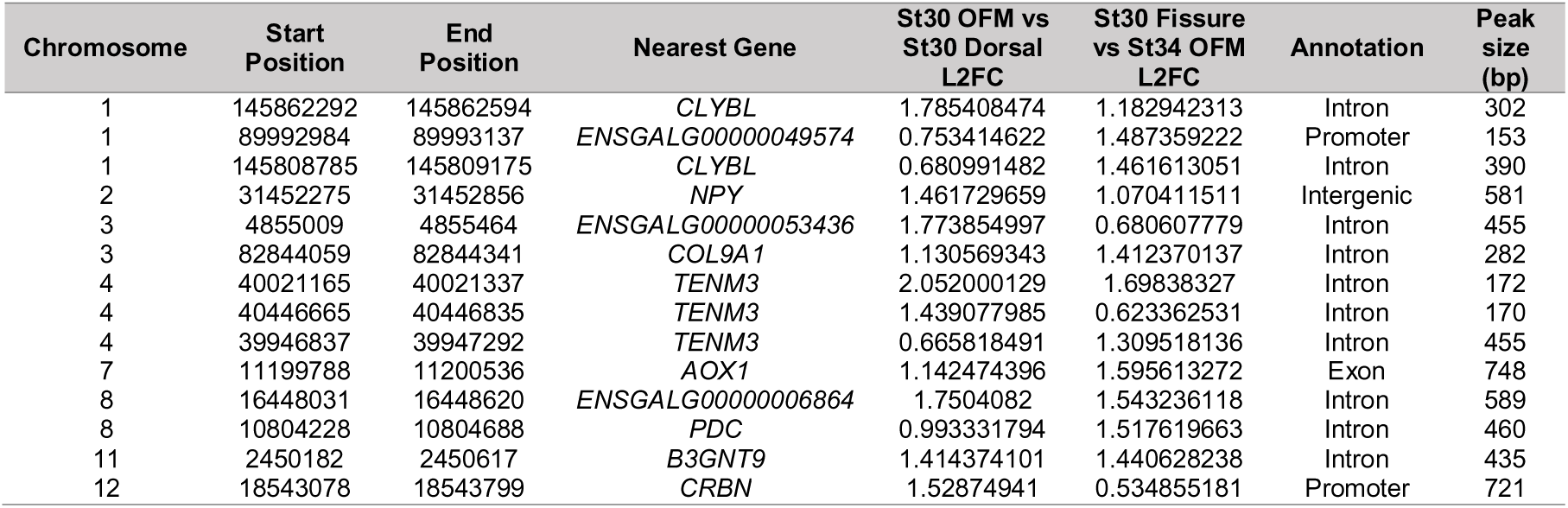
Top 10 genes associated with highest DA fusion specific proximal peak coordinates. Adjusted *P*-value < 0.05.

### Identifying candidate transcription factor networks in the developing eye

We then asked if our data could provide evidence for specific TF activity to infer gene regulatory networks during OFC. We first performed unbiased TF binding motif identification from our whole genome ATAC-seq peaks using HOMER *de novo* motif enrichment analyses. Motifs of 8-12bp in length were searched for, representing the typical lengths of TF binding motifs^70^. Enrichment was determined by comparing the appearance of motifs within target sequences (e.g. fusion enriched peaks) versus background. We identified 4 TF motifs associated with fusion, and 3 TF motifs associated with OFM enriched peaks (*P* <1×10^-50^) (**Figure 3A-B**). Notably, enrichment for the TEAD family motif (TEAD3; (*P* < 1×10^-156^) was unique to the fusion specific peaks, while motifs corresponding to the homeobox proteins LHX2/VSX1/VAX2 and the fusion specific DEG ALX1 (which all have largely identical motifs to each other - **Supplemental Figure S4A**), were identified in the fusion and OFM enriched peaks, as well as SOX and ZIC family TF motifs. In contrast, these motifs were not identified in in either the dorsal enriched or fusion repressed peak sets (**Figure 3C-D**), but these comparison groups both included significant enrichment for retinoic-acid-related orphan receptor (ROR) transcription factor subfamily motifs (*P* < 1×10^-100^). At the transcriptional level, *ALX1*, *ZIC1*, *LHX2* and *SOX5* were all enriched in the OFM, whereas *RORB* expression was enriched at HH34 and dorsally at HH30, while *VAX2* and *VSX1* expression were both dorsally enriched (**Supplemental Figures S4C, S2C**).

**Figure 3.**
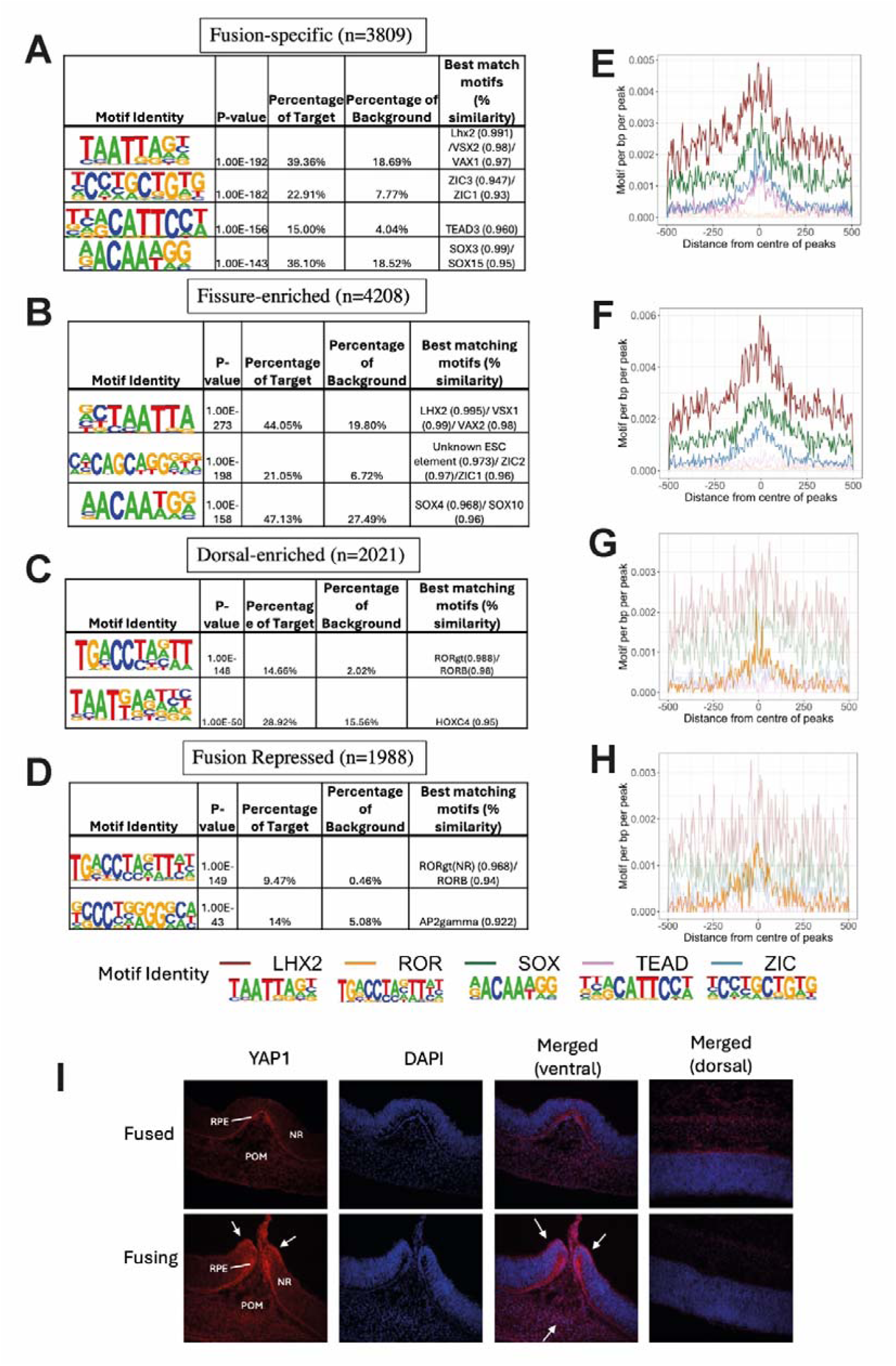
Transcription factor motif enrichment in OFC. (**A-D**) Number of peaks and highest match *de novo* motifs identified by HOMER for each context specific category. Only motifs with a percentage of target to background ratio > 1.5 were included. (**E-H**) Histograms showing motif occurrence distribution around the centre of peaks corresponding to analysis in A-D. Lines without observable enrichment are translucent. Motif consensus sequences and their likely TF identities are shown below and heights correlate to the frequency of base occurrence. (**I**) YAP1 immunofluorescence during chicken OFC at HH30 showed highest signal (arrows) in the edges of the fusing OFM and in the fused margins. In contrast, low signal was detected in dorsal tissue. *Abbreviations*: RPE, retinal pigmented epithelium, NR, neural retina; POM, periocular mesenchyme.

### TF motif analysis predicts TEAD activity in OFC

We then examined the motif density observed for these TFs in 1000 bp windows (± 500bp of peak centre) in all sample peaks. Enrichment of ZIC, SOX, TEAD and LHX2 motifs was observed to spike near the centre of the fusion specific peaks, while ROR showed no observable change for these peaks and had a low motif density (**Figure 3E**). In the OFM enriched data, ZIC, SOX, and LHX2 motifs were enriched at the centre of peaks, but not TEAD or ROR (**Figure 3F**). In contrast, the ROR motif was enriched at the centre of peaks within both fusion repressed and the dorsal enriched peak set, while TEAD, SOX, VAX/LHX2/VSX2 and ZIC motifs were not enriched at the centre (**Figure 3G-H**). We then analysed expression of *TEAD* and *ROR* in the RNA-seq data across the samples and found that none of the *TEAD* paralogues (*TEAD1*, *TEAD3*, *or TEAD4*) were enriched for expression in the HH30 OFM, however all were generally expressed at a higher level at HH30 compared to HH34 (**Supplemental Figure S4B**). The TEAD transcriptional co-activator transcription factor *YAP1* was also most highly expressed at HH30 compared to HH34 but was not specific to the OFM. However, immunofluorescence analysis at HH30 showed apparent enrichment for YAP1 at the protein level in retina-RPE and at the edges of the fissure margins in the HH30 OFM compared to dorsal eye NR and RPE regions at this stage (**Figure 3I**). In combination, these data indicate a specific involvement for YAP-TEAD activity during fusion of the OFM.

### TF foot-printing analysis indicates fusion specific regulatory networks

We then aimed to further support this data by determining TFBS occupancy dynamics using foot-printing analysis^44^, where direct TF binding sterically inhibits Tn5 transposition within otherwise accessible chromatin. In terms of overall binding differences, multiple *de novo* motifs with predicted TEAD binding were preferentially bound to genome-wide HH.30 associated OFM peaks when compared with peaks enriched in both HH.30 dorsal and HH34 fissure (**Figure 4A**). In contrast, motifs with predicted ROR binding showed preferential occupancy in dorsal peaks, or in HH34 OFM, further indicating fusion specific repression. To investigate the *bona fide* binding occupancy of TEAD motifs to peaks with fusion specific accessibility, we then examined the aggregated signal at the TEAD motif sites of all 105 fusion enriched peaks. An observable footprint, with a depression of signal in the centre of a generally increased accessible region was most obvious for TEAD in the HH.30 fissure (**Figure 4B**). In comparison, the TEAD motif in the dorsal HH30 peaks did not show any clear footprint, whereas there was clearly observable footprinting for ROR motifs. To illustrate the combined influence of TF binding at fusion, we analysed chromatin at the single gene level. *CLYBL* illustrated fusion specific gene expression from our RNA-seq analysis (**Figure 2A & C**). We first confirmed the expression was indeed fusion specific using *in situ* hybridisation. We found strong expression in the open fissure but none in the dorsal retina or in the nascently fused fissure (**Figure 4C**). Thus, *CLYBL* is a gene with highly specific spatial and temporal expression during OFC and therefore is likely to have high degree of dynamic transcriptional regulation. We therefore looked at the chromatin regulation at the *CLYBL* locus and found 2 clear intronic peaks with fusion specific enrichment containing motifs for *TEAD*, *SOX5* and *ZIC1* (**Figure 4D**). In combination, *de novo* motif enrichment and DNA foot-printing analyses suggest that the regulatory landscape is markedly different between the fissure and dorsal tissue during OFC, and that TEAD, SOX, ZIC and homeobox TF activities are likely to be important regulators of fusion specific gene expression.

**Figure 4.**
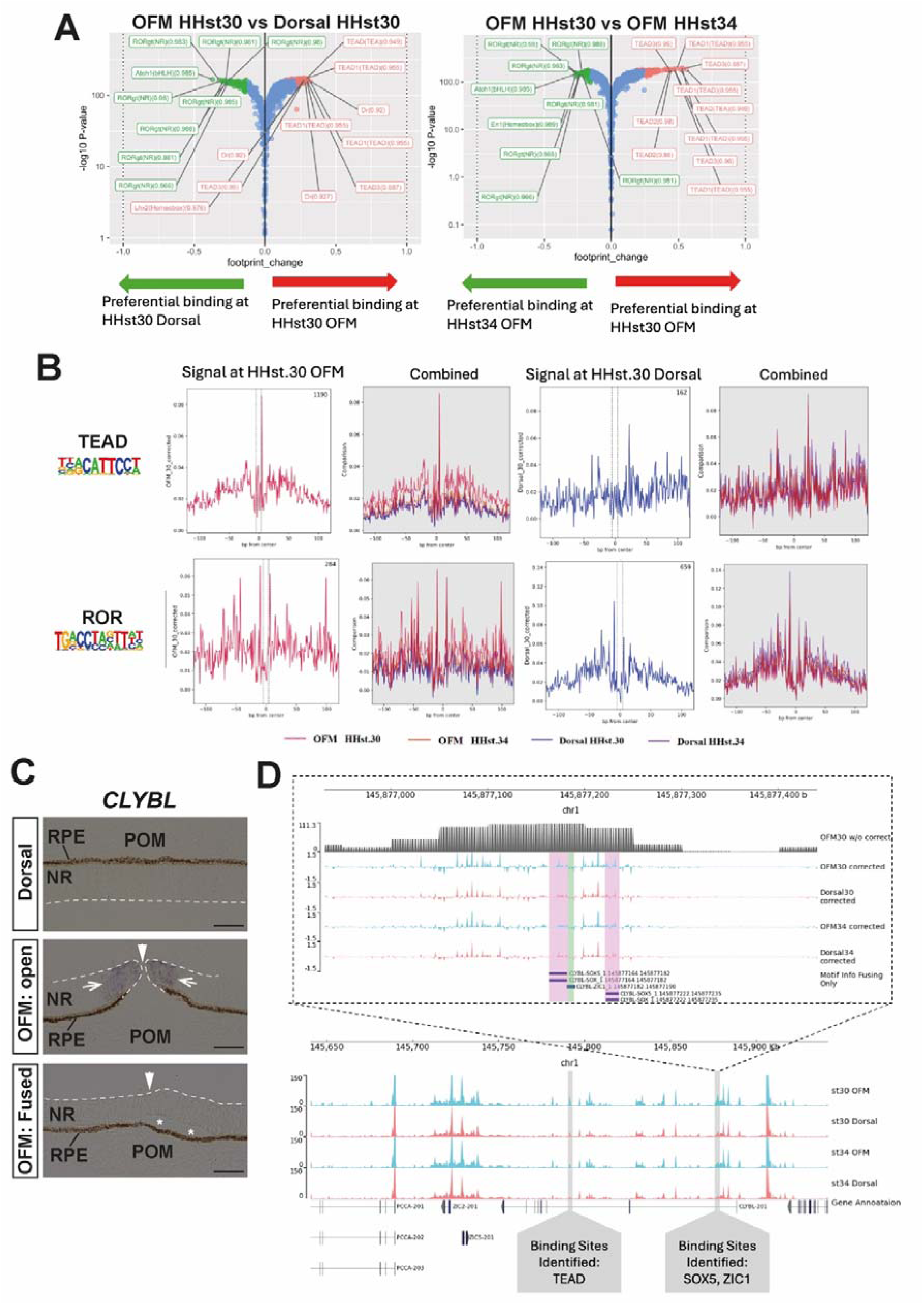
ATAC-seq foot-printing shows differential activities for TEAD and ROR motifs. (**A**) Volcano plots showing the binding preference of each HOMER de novo motif with comparisons between HH30 fissure to HH30 dorsal (*left*), and HH.30 fissure to HH.34 fissure (*right*). Red and green dots highlighted the TF motifs which have a top or bottom 5^th^ percentile in terms of OFM binding, respectively. The top 10 TF motifs with the strongest differential binding are labelled, together with their percentage identity. (**B**) Aggregated plots showing the *bona fide* footprint of TEAD and ROR motifs (shown). *Left*: combined signals of all TF binding occurrence sites within fusion specific enriched peaks. *Right*: Combined signals of all TF binding occurrence sites within fusion repressed peaks. (**C**) Brightfield microscopy images of *in situ* hybridisation of *CLYBL* (purple stain) in sagittal sections in the HH30 chicken eye. Fusion specific expression was observed in cells at the edges of the open OFM (open arrows), with no detectable expression in the midline of the nascently fused OFM, or in the dorsal eye (*n* = 3). (**D**) Screenshot of the *CLYBL* locus in the chicken genome showing ATAC-seq tracks for each of the 4 conditions studied and the genome annotation for *CLYBL*. HH30 fusion specific (HH30 only enriched) peaks are highlighted, and an enlarged image of the peak containing ZIC1/SOX5 TF motifs is presented, alongside a second fusion specific TEAD motif containing peak.

### Identification of orthologous human loci from chicken DA peaks

Finally, we asked whether the gene regulatory dynamics we observed in the chicken eye had relevance to human OFC. We took the 105 peaks with the most differential accessibility in proximity (40kb + gene length) to fusion-specific DEGs, together with OFM enriched DA peaks, and DA peaks representing the top 10% DA loci for fusion genome-wide (i.e. that were not associated with fusion-specific DEGs) to perform DNA sequence similarity local alignment searches (BLASTn) of these peak sequences against the human genome to identify orthologous regions **(Supplemental Figure 5)**^71^. Using this approach, we identified 33 loci in humans with sequence identity >80% and BLAST E-value < 1 x 10^-50^ (**Supplemental Table S4**). Among these, 29 loci overlapped with at least one ENCODE-annotated *cis*-regulatory regions (**Supplemental Table S4**), representing a significant enrichment for mapping to CREs compared with GC-matched background regions (P = 1.24 x 10^-4^). The closest neighbouring genes of these orthologous regions were identical between chicken and human. Also, we identified orthologous regions in proximity to the chicken fusion-specific DEGs *CLYBL*, *AOX1*, *BMP5*, *TES*, and *FLRT3*, or genes that were already associated with colobomas in humans or animal models, such as *PAX2*, *CHRDL1*, *WNT7B, VAX1,* and *BMPR1B*^10,72^. This data suggests these regions may regulate the expression of genes important for optic fissure closure across a range of species and represents the first evidence for *cis* regulation in this biological process.

## DISCUSSION

As few patients with coloboma currently receive a genetic diagnosis^1–3^, this study provides novel genetic resources to help support the identification, interrogation, and interpretation of non-coding variants in human coloboma and MAC patients. As other ATACseq datasets have recently become available for human optic vesicle cultures and retinal development out-with the optic fissure, our work adds to growing list of resources that can be exploited to delineate the activities and requirements for specific non-coding loci in a range of eye developmental contexts. Given the low global incidence of coloboma and the low rate of genes identified with recurrently identified coding-region mutations in unrelated patients, it is likely that the identification of causative variants in non-coding regions will be challenging. However, the increased application of whole genome sequencing for many patients and family members globally, and the growing collections and curation of sequencing data for rare disease cohorts (including MAC patients), has generated large genomic resources that can now be interrogated for SNPs, indels, and copy number variants in non-coding regions in patient sequences, where our data provides supportive evidence for likely functionality in OFC and eye development.

We also identified compelling evidence for TEAD as a transcriptional regulator for OFC. TEAD is a TF that is constitutively located in the nucleus that only becomes transcriptionally active when bound to nuclear translocated YAP1 protein^73^. TEAD/YAP signalling has previously been linked to a role in broader eye development by regulating the differentiation of neural progenitors into RPE or neural retina fates, and *in vivo YAP1* loss of function mutational studies have observed reduced RPE with colobomas^74^. Fusion at the OFM is directly mediated through EMT-like behaviours of a discrete population of “pioneer cells” that are physically located at the edges of the optic fissure margins at the junction between RPE or neural retina cell types and which show characteristics of both^52,75,76^. Therefore, disruption to this signalling pathway may affect the development or maintenance of pioneer cells, or their normal functions of mediating fusion in OFC. We showed enrichment for TEAD motifs and TF occupancy in our fusion specific peaks and observed increased YAP1 protein localisation specifically at the edges of the OFM. Heterozygous missense mutations in *YAP1* cause coloboma and microphthalmia in humans^77,78^, however, no causative mutations in *TEAD* genes have yet been discovered. Our data suggests that this is likely, and that TEAD genes should be included in genetic screens in MAC patients, but also that revealing YAP1/TEAD transcriptional target genes and their regulatory regions could reveal further coloboma candidates.

We also detected both motif enrichment and preferential binding of ROR family transcription factors specifically in fusion-repressed peaks. In line with the ATAC-seq occupancy data, *RORB* showed repressed gene expression specifically in the actively fusing optic fissure. *RORB* was previously shown to have dorsally enriched expression in developing mouse eyes, and loss of function for *RORB* caused microphthalmia in knockout mice^79^. RORB ligands includes all-*trans* retinoic acid, a metabolic product of aldehyde dehydrogenase encoded by *ALDH1A3*, which displays fusion specific expression in the developing vertebrate eye^80,81^. Since all-*trans* retinoic acid inhibits the transcription factor activity of RORB^80^, this suggests ALDH1A3 may inhibit RORB activity and its downstream pathways to maintain dorsal-ventral patterning in the optic cup and regulation of OFC.

Although this study was able to detect candidate regulatory loci, a main limitation was that we did not perform any *in vivo* testing of their enhancer activity. However, now these regulatory elements have been identified, subsequent work can be focused on determining their functional role in eye development and confirming their regulatory target genes. The chicken is a versatile model organism for performing *in vivo* enhancer reporter assays in early development^23,24,82^, and enhancers with eye-specific activity have previously been successfully tested for functionality within the chicken eye^83^. As more MAC and coloboma associated intragenic variants are identified in human patients, robust and scalable approaches to determine causality will be required. A recent study deleted conserved non-coding elements in *xenopus* embryos where the orthologous locus was found with a variant in MAC patients and showed that the edited allele resulted in reduced expression in the proximal *MAB21L2* gene^84^. In addition, enhancer testing in zebrafish is well established and can detect functional differences between allele variants in developing embryos^85^. Recently, chicken embryos with constitutive expression of CAS9 have also been generated^86^ and could provide a powerful tool for testing of regulatory elements through gene editing, where the presence or absence of a coloboma or small eye could phenotypically confirm a functional requirement for the edited candidate locus.

We applied sequence homology searching to identify regions in the human genome orthologous to the peaks identified in chicken and found 29 proximal to known OFC or coloboma genes. This approach stringently selected conserved human sequences with high sequence-level identity, however some functionally conserved enhancers are not conserved at the DNA sequence level^87^, and therefore our approach may have missed sequence degenerate loci that regulate OFC. An alternative approach may be to search using TFBS combinations and synteny, including the arrangement, orientation and order of TFBS. Our study empowers future work in this area as we identified a range of transcription factors which either exhibited occupancy bias or which showed enrichment for their TFBS sequences in the fusion specific and fusion enriched peaks, such as homeodomain TFBS binding motifs including *VAX1/2* and *LHX2*^63,88^, which were also highly expressed TFs in the fusing optic fissure.

In conclusion, this study leverages conserved vertebrate eye development in the chicken embryo to deliver the first evidence for dynamic *cis* regulatory activity of gene expression during OFC, and identifies specific genomic loci and putative GRNs for future mutational and activity analyses to determine their precise roles in this developmental context and improve the genetic diagnosis of human colobomas.

## DECLARATIONS

### Ethics approval and consent to participate

Only chicken embryos at non-protected embryonic stages under the Animals (Scientific Procedures) Act 1986 were used for this study and no regulated procedures were performed. JR holds a Home Office Project Licence PPL0018931. Human data was not used in this study.

### Consent for publication

Not applicable.

### Author Contributions

Conceptualization, J.P. and J.R.; methodology, B.H.C.C., J.P. and J.R.; software, B.H.C.C., J.P.; Training and supervision, H.H.; investigation, B.H.C.C. and J.R.; formal analysis, B.H.C.C., and J.R.; resources, M.M. and J.R.; data curation, B.H.C.C., ; writing—original draft preparation, B.H.C.C, and J.R.; writing—review and editing, B.H.C.C., H.H. J.R. J.P. and M.M.; funding acquisition, J.R. and M.M. All authors have read and agreed to the published version of the manuscript.

### Data Availability Statement

The dataset(s) supporting the conclusions of this article are available in the NCBI Gene Expression Omnibus database (http://www.ncbi.nlm.nih.gov/geo) with accession number GSE318644. All custom codes are available freely upon request.

## Acknowledgments

We would like to thank the staff at the Greenwood Building (National Avian Research Facility, Roslin Institute) for chicken husbandry, and Anna Raper at the Bioimaging and Flow Cytometry Facility (Roslin Institute) for cell sorting. We are also very grateful to Tatjana Sauka-Spengler and Ruth Williams for help and advice with ATAC-seq preparations in chicken embryos, and Andrew Papanastasiou and Jessica Powell for helpful discussions over ATAC-seq analyses.

## SUPPLEMENTAL FIGURES

**Figure S1.**
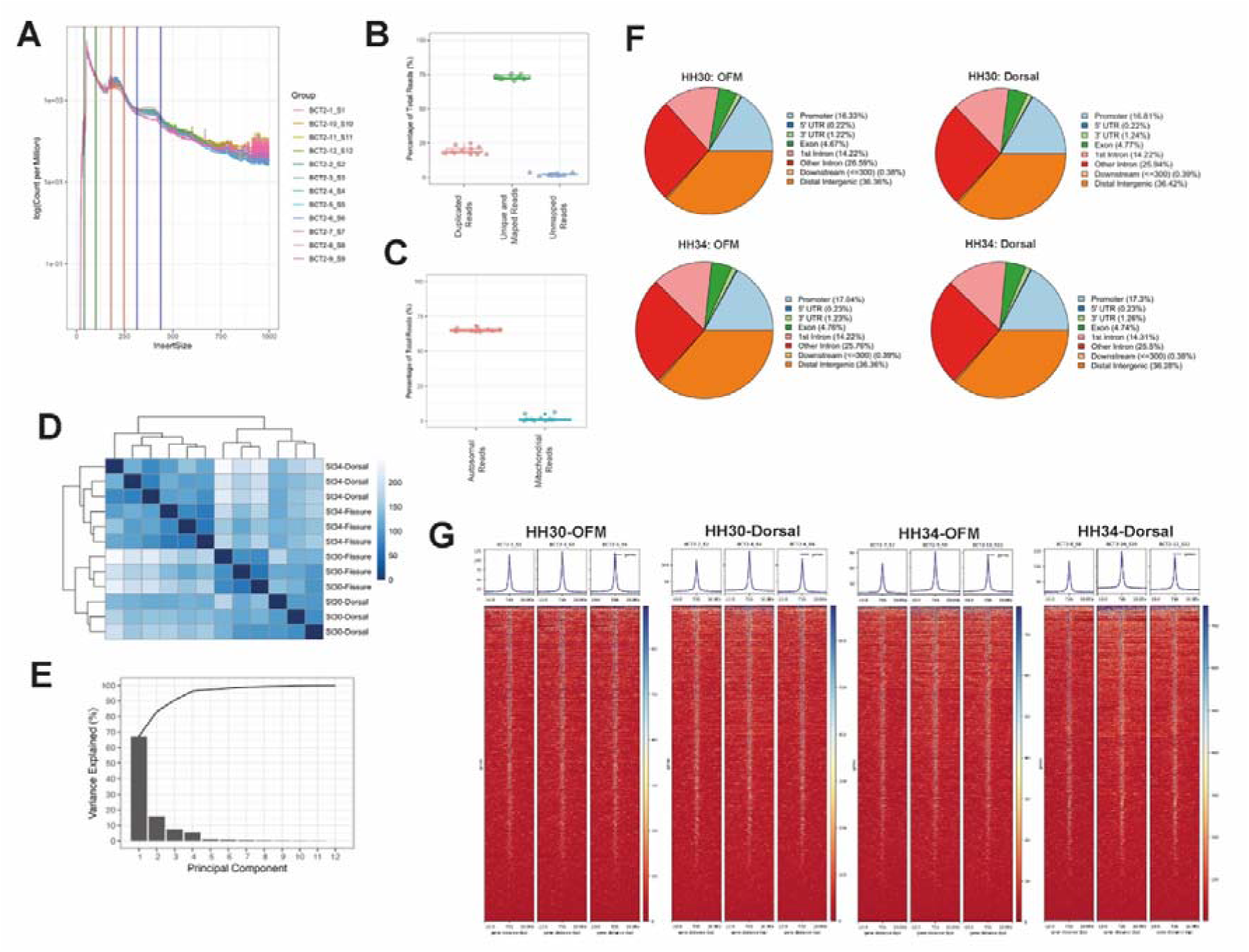
Profiling for open chromatin during optic fissure closure. (**A**) Nucleosomal profiling matrix for ATAC-seq dataset. Green, red and blue vertical-line bounded regions correspond to free DNA fragments, mono-nucleosomal, and di-nucleosomal fragments, respectively. (**B**-**C**) Metrics for duplicated and mapped reads (**B**), and mitochondrial versus autosomal mapped reads (**C**). (**D**) Euclidean distance plot showing sample to sample difference of reads mapping to peaks. (**E**) Scree plot showing the percentage of sample-to-sample variance explained by different Principal Components. (**F**) Distribution of genomic annotations of all peaks at each condition. (**G**) TSS enrichment plot and heatmaps of open chromatin regions for each sample replicate across the four conditions. Colour indicates normalised number of reads per bin for each condition, with red and blue to indicate low and high reads per bin, respectively. TSS is labeled as 0 on the x-axis. See **Supplemental Table S1** for sample annotation identifiers.

**Figure S2.**
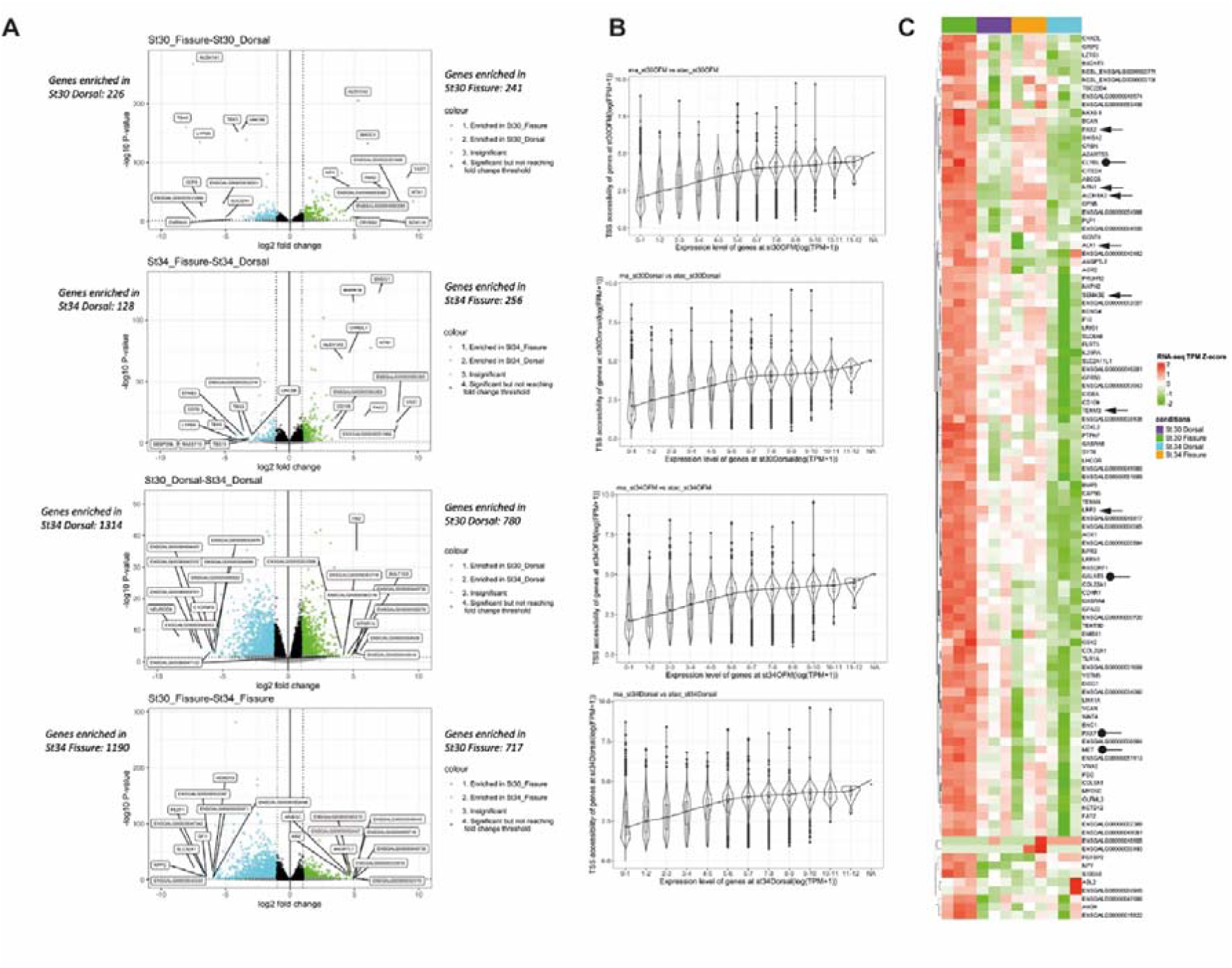
Chicken OFC RNA-seq analysis and correlation with ATAC-seq. (**A**) Volcano plots showing differentially expressed genes for each comparison. The top 10 DEGs in both conditions of the pairwise comparison are labelled on the graph. Dotted vertical lines indicate L_2_FC of >1 and <-1; horizontal dotted line indicates adj-*P* 0.05 threshold. (**B**) Violin plots showing distributions of transcription start site accessibilities of binned groups of coding genes with expression levels ranging from a log_10_(1+Transcript per million) value of 0-1 to 11-12 in HH.30 fissure, HH.30 dorsal, HH.34 fissure, and HH.34 dorsal. (**C**) Heatmap showing z-scores of TPMs for HH30 OFM specific 106 DEGs compared to the other spatial and temporal categories. Arrows indicate known human coloboma genes, round-ended lines are genes with verified fusion specific expression.

**Figure S3.**
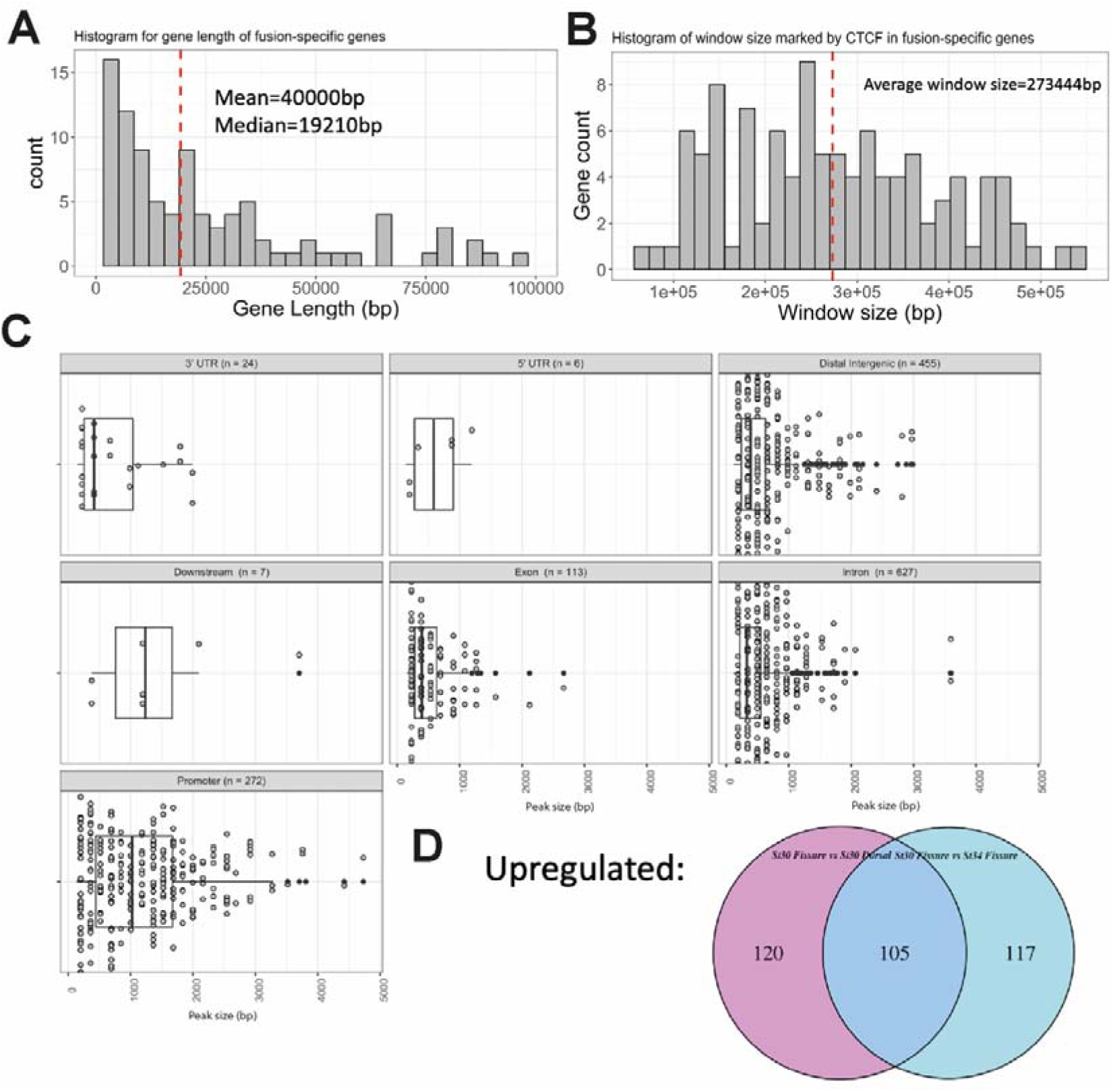
Fusion specific DA chromatin loci analysis. (**A**) Analysis of all 106 fusion enriched DEGs revealed a mean gene length of 40kb, with 9 genes exceeding 100kb. (**B**) Dot plot showing the distribution of peak lengths for all fusion specific proximal peaks belonging to each category of genomic annotation. The number of peaks belonging to each category are indicated and X-axis corresponds to peak size in bp. (**C**) Histogram showing the distribution of the CTCF-defined window size for each fusion specific enriched gene. The highest CTCF signal within 250bp upstream of transcription start site (TSS) or downstream of the gene marked the predicted boundary respectively. The red dotted line indicates the average window size across all fusion specific enriched genes. 5 of the 106 genes contained no CTCF binding signal either upstream or downstream within the 250kb window and the search window was extended to 500kb upstream and downstream for these genes. (**D**) Venn diagram indicating fusion specific DA peaks determined by comparison between HH30 OFM versus HH30 dorsal and HH34 OFM DA peaks.

**Figure S4.**
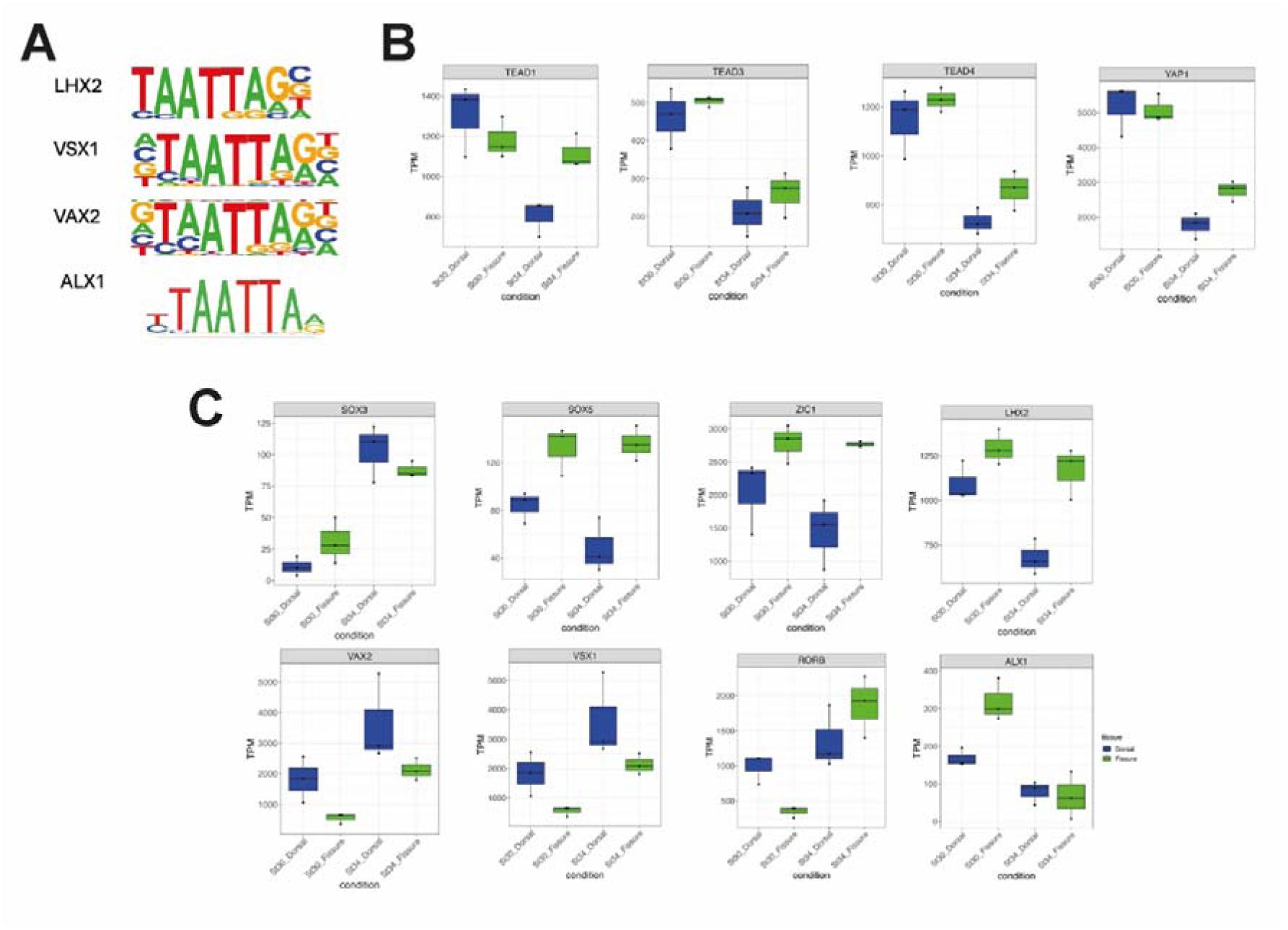
Transcription factors in the chicken optic fissure. **(A)** Shared sequence similarities for homeodomain motifs LHX2, VAX1, ALX1 and LHX2. (**B**) TPM plot showing expression patterns of *TEAD1*, *TEAD3*, *TEAD4* and *YAP1* from the RNA-seq analysis**. (C)** TPM of transcription factors expression levels from the RNA-seq data. Only values for transcription factor genes with a motif enrichment indicated by HOMER were plotted.

**Figure S5.**
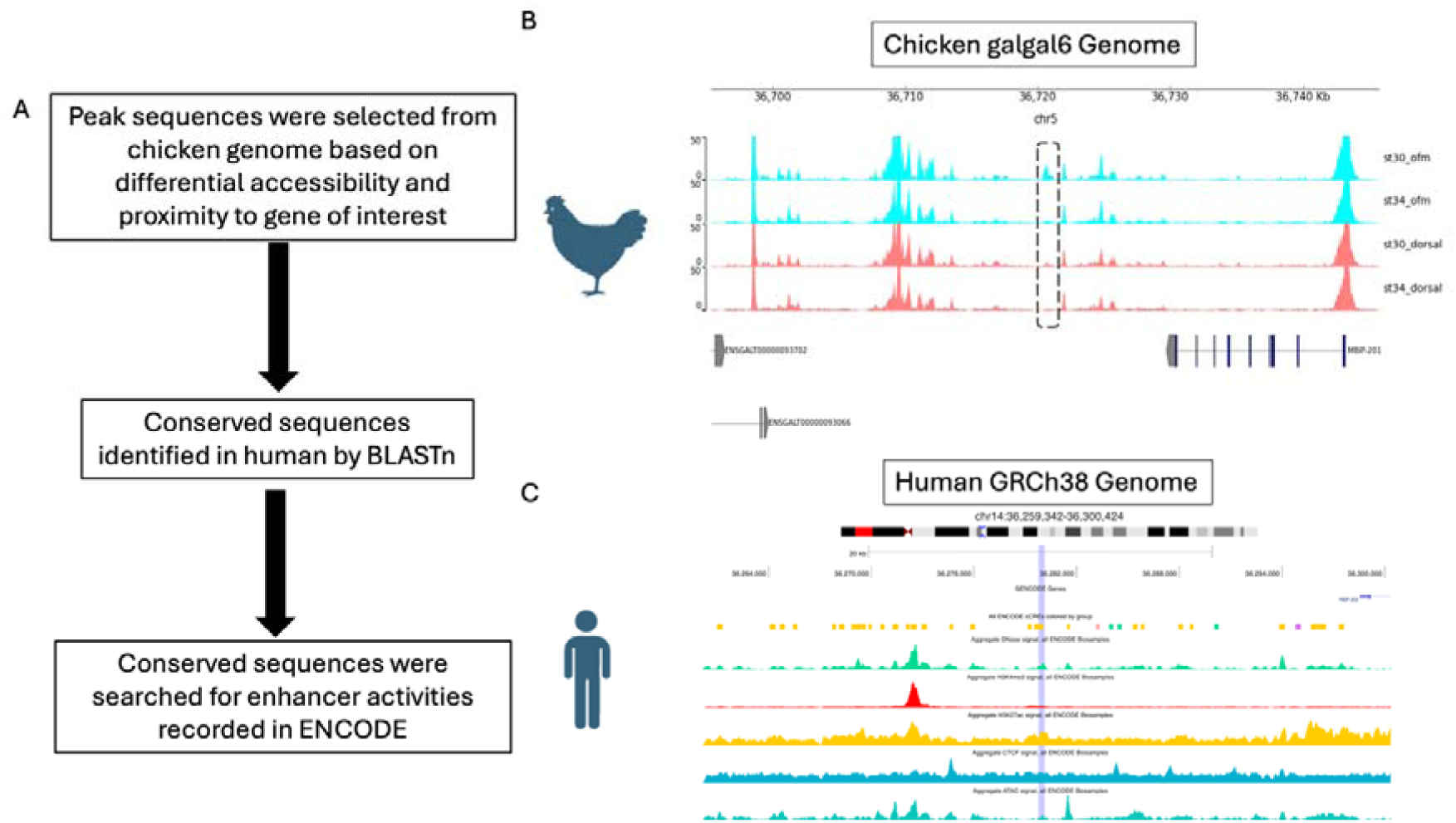
BLAST-based lifting of chicken putative enhancer coordinates to identify human enhancer regions. (**A**) Workflow illustrating the process of lifting chicken putative OFC enhancer coordinates from the *Gallus gallus* (galGal6) genome to the human GRCh38 genome using a BLAST-based approach. (**B**) Genomic view of the galGal6 chicken genome at chr5:36,695,374–36,745,738. Annotated transcripts within the region are shown at the bottom. The hatched box highlights the position of a representative differentially accessible peak. (**C**) Corresponding genomic view in the human GRCh38 genome at chr14:36,245,298–36,314,609. ENCODE-defined enhancer regions are shown in yellow. The purple box indicates BLAST-detected region of high sequence homology which mapped to an ENCODE enhancer. This diagram was prepared using SCREEN Registry V4^89^.

## REFERENCES

1. Williamson, K. A. & FitzPatrick, D. R. The genetic architecture of microphthalmia, anophthalmia and coloboma. Eur. J. Med. Genet. 57, 369–380 (2014).

2. Patel, A. & Sowden, J. C. Genes and pathways in optic fissure closure. Semin. Cell Dev. Biol. 10.1016/j.semcdb.2017.10.010 doi:10.1016/j.semcdb.2017.10.010.

3. Harding, P. et al. Real-world clinical and molecular management of 50 prospective patients with microphthalmia, anophthalmia and/or ocular coloboma. British Journal of Ophthalmology bjo-2022-321991 (2022) doi:10.1136/bjo-2022-321991.

4. Selzer, E. B. et al. Review article Review of evidence for environmental causes of uveal coloboma. Surv. Ophthalmol. 67, 1031–1047 (2021).

5. Chan, B. H. C., Moosajee, M. & Rainger, J. Closing the Gap: Mechanisms of Epithelial Fusion During Optic Fissure Closure. Frontiers in Cell and Developmental Biology vol. 8 Preprint at 10.3389/fcell.2020.620774 (2021).

6. Morrison, D. et al. National study of microphthalmia, anophthalmia, and coloboma (MAC) in Scotland: investigation of genetic aetiology. J. Med. Genet. 39, 16–22 (2002).

7. Shah, S. P. et al. Anophthalmos, microphthalmos, and coloboma in the United Kingdom: Clinical features, results of investigations, and early management. Ophthalmology 119, 362–368 (2012).

8. Patel, A., Anderson, G., Galea, G., Balys, M. & Sowden, J. C. A molecular and cellular analysis of human embryonic optic fissure closure related to the eye malformation coloboma. Development 44, dev.193649 (2020).

9. Trejo-Reveles, V. et al. Identification of Novel Coloboma Candidate Genes through Conserved Gene Expression Analyses across Four Vertebrate Species. Biomolecules vol. 13 Preprint at 10.3390/biom13020293 (2023).

10. Owen, N. et al. Identification of 4 novel human ocular coloboma genes ANK3, BMPR1B, PDGFRA, and CDH4 through evolutionary conserved vertebrate gene analysis. Genetics in Medicine 24, 1073–1084 (2022).

11. Chang, L., Blain, D., Bertuzzi, S. & Brooks, B. P. Uveal coloboma: clinical and basic science update. Curr. Opin. Ophthalmol. 17, 447–470 (2006).

12. Harding, P. et al. Variant-specific disruption to notch signalling in PAX6 microphthalmia and aniridia patient-derived hiPSC optic cup-like organoids. Biochimica et Biophysica Acta (BBA) - Molecular Basis of Disease 1871, 167869 (2025).

13. Lauderdale, J. D., Wilensky, J. S., Oliver, E. R., Walton, D. S. & Glaser, T. 3’ deletions cause aniridia by preventing PAX6 gene expression. Proc. Natl. Acad. Sci. U. S. A. 97, 13755–9 (2000).

14. Hall, H. N. et al. Short-read whole genome sequencing identifies causative variants in most individuals with previously unexplained aniridia. J. Med. Genet. 61, 250–261 (2023).

15. Bhatia, S. et al. Disruption of autoregulatory feedback by a mutation in a remote, ultraconserved PAX6 enhancer causes aniridia. Am. J. Hum. Genet. 93, 1126–34 (2013).

16. Mitchell, L. A. et al. Axenfeld-Rieger syndrome associated with a megabase-scale inversion separating PITX2 from a conserved enhancer locus. medRxiv 10.1101/2025.06.05.25327661 (2025) doi:10.1101/2025.06.05.25327661.

17. Wormser, O. et al. IHH enhancer variant within neighboring NHEJ1 intron causes microphthalmia anophthalmia and coloboma. NPJ Genom. Med. 8, 22 (2023).

18. Ceroni, F. et al. Deletion upstream of MAB21L2 highlights the importance of evolutionarily conserved non-coding sequences for eye development. Nat. Commun. 15, 9245 (2024).

19. Bernstein, C. S. et al. The cellular bases of choroid fissure formation and closure. Dev. Biol. 440, 137–151 (2018).

20. Hardy, H. et al. Detailed analysis of chick optic fissure closure reveals Netrin-1 as an essential mediator of epithelial fusion. Elife 8, (2019).

21. Hardy, H. & Rainger, J. Cell adhesion marker expression dynamics during fusion of the optic fissure. 10.1101/2023.08.01.551419 https://doi.org/10.1101/2023.08.01.551419 (2023).

22. Chan, B. H. C. et al. A stable NTN1 fluorescent reporter chicken reveals cell specific molecular signatures during optic fissure closure. Sci. Rep. 15, 10096 (2025).

23. Smith, E. L., Mok, G. F. & Münsterberg, A. Investigating chromatin accessibility during development and differentiation by ATAC-sequencing to guide the identification of cis-regulatory elements. Biochem. Soc. Trans. 50, 1167–1177 (2022).

24. Williams, R. M. et al. Reconstruction of the Global Neural Crest Gene Regulatory Network In Vivo. Dev. Cell 51, 255–276.e7 (2019).

25. Mok, G. F. et al. Characterising open chromatin in chick embryos identifies cis-regulatory elements important for paraxial mesoderm formation and axis extension. Nat. Commun. 12, 1157 (2021).

26. Zhang, X. et al. ATAC-seq and RNA-seq analysis unravel the mechanism of sex differentiation and infertility in sex reversal chicken. Epigenetics Chromatin 16, 1–17 (2023).

27. Li, J. et al. Single-nucleus transcriptional and chromatin accessible profiles reveal critical cell types and molecular architecture underlying chicken sex determination. J. Adv. Res. 70, 29–43 (2025).

28. Li, J. et al. Time-Course Transcriptional and Chromatin Accessibility Profiling Reveals Genes Associated With Asymmetrical Gonadal Development in Chicken Embryos. Front. Cell Dev. Biol. 10, 1–12 (2022).

29. Hamburger, V. & Hamilton, H. L. A series of normal stages in the development of the chick embryo. J. Morphol. 88, 49–92 (1951).

30. Hamburger, V. & Hamilton, H. L. A series of normal stages in the development of the chick embryo. 1951. Dev. Dyn. 195, 231–272 (1992).

31. Williams, R. M. & Sauka-Spengler, T. Dissociation of chick embryonic tissue for FACS and preparation of isolated cells for genome-wide downstream assays. STAR Protoc. 2, 100414 (2021).

32. Corces, M. R. et al. An improved ATAC-seq protocol reduces background and enables interrogation of frozen tissues. Nat. Methods 14, 959–962 (2017).

33. Martin, M. Cutadapt removes adapter sequences from high-throughput sequencing reads. Molecular Biology Network 17, 10–12 (2011).

34. Langmead, B., Trapnell, C., Pop, M. & Salzberg, S. L. Ultrafast and memory-efficient alignment of short DNA sequences to the human genome. Genome Biol. 10, R25 (2009).

35. Orchard, P., Kyono, Y., Hensley, J., Kitzman, J. O. & Parker, S. C. J. Quantification, Dynamic Visualization, and Validation of Bias in ATAC-Seq Data with ataqv. Cell Syst. 10, 298–306.e4 (2020).

36. Li H, Handsaker B, Wysoker A, Fennell T, Ruan J, Homer N, Marth G, Abecasis G, Durbin R, 1000 Genome Project Data Processing Subgroup. The Sequence Alignment/Map format and SAMtools. Bioinformatics 25, 2078–9 (2009).

37. Gaspar, J. M. Improved peak-calling with MACS2. Preprint at 10.1101/496521 (2018).

38. Lopez-Delisle, L. et al. pyGenomeTracks: reproducible plots for multivariate genomic datasets . Bioinformatics 37, 422–423 (2021).

39. Quinlan, A. R. & Hall, I. M. BEDTools: a flexible suite of utilities for comparing genomic features. Bioinformatics 26, 841–2 (2010).

40. Liao, Y., Smyth, G. K. & Shi, W. featureCounts: an efficient general purpose program for assigning sequence reads to genomic features. Bioinformatics 30, 923–930 (2014).

41. Love, M. I., Huber, W., & Anders, S. Moderated estimation of fold change and dispersion for RNA-seq data with DESeq2. Genome Biol. 15, 1–21 (2014).

42. Lawrence, M., et al. Software for Computing and Annotating Genomic Ranges. PLoS Comput. Biol. 9, e1003118 (2013).

43. Yu, G., Wang, L.-G. & He, Q.-Y. ChIPseeker: an R/Bioconductor package for ChIP peak annotation, comparison and visualization. Bioinformatics 31, 2382–2383 (2015).

44. Heinz, S. et al. Simple Combinations of Lineage-Determining Transcription Factors Prime cis-Regulatory Elements Required for Macrophage and B Cell Identities. Mol. Cell 38, 576–589 (2010).

45. Bentsen, M. et al. ATAC-seq footprinting unravels kinetics of transcription factor binding during zygotic genome activation. Nat. Commun. 11, 4267 (2020).

46. Weirauch, M. T. et al. Determination and Inference of Eukaryotic Transcription Factor Sequence Specificity. Cell 158, 1431–1443 (2014).

47. Rauluseviciute, I., et al. JASPAR 2024: 20th anniversary of the open-access database of transcription factor binding profiles. Nucleic Acids Res. 52, D174–D182 (2024).

48. Kadota, M. et al. CTCF binding landscape in jawless fish with reference to Hox cluster evolution. Sci. Rep. 7, (2017).

49. Trevers, K. E. et al. A gene regulatory network for neural induction. Elife 12, (2023).

50. Camacho, C., et al. BLAST+: architecture and applications. BMC Bioinformatics 10, 421 (2009).

51. Abascal, F. et al. Expanded encyclopaedias of DNA elements in the human and mouse genomes. Nature 583, 699–710 (2020).

52. Ho, B. et al. A stable NTN1 fluorescent reporter chicken reveals cell specific molecular signatures during optic fissure closure. 1–12 (2025).

53. Richardson, R. et al. Transcriptome profiling of zebrafish optic fissure fusion. Sci. Rep. 9, 1–12 (2019).

54. Brown, J. D. et al. Expression profiling during ocular development identifies 2 Nlz genes with a critical role in optic fissure closure. Proc. Natl. Acad. Sci. U. S. A. 106, 1462–7 (2009).

55. Dorgau, B. et al. Single-cell analyses reveal transient retinal progenitor cells in the ciliary margin of developing human retina. Nat. Commun. 15, 1–17 (2024).

56. Buenrostro, J. D., Wu, B., Chang, H. Y. & Greenleaf, W. J. ATAC-seq: A Method for Assaying Chromatin Accessibility Genome-Wide. Curr. Protoc. Mol. Biol. 1–10 (2016) doi:10.1002/0471142727.mb2129s109.ATAC-seq.

57. Orchard, P., Kyono, Y., Hensley, J., Kitzman, J. O. & Parker, S. C. J. Quantification, Dynamic Visualization, and Validation of Bias in ATAC-Seq Data with ataqv. Cell Syst. 10, 298–306.e4 (2020).

58. Akoglu, H. User’s guide to correlation coefficients. Turk. J. Emerg. Med. 18, 91–93 (2018).

59. Sureshchandra, S. et al. Chronic heavy drinking drives distinct transcriptional and epigenetic changes in splenic macrophages. EBioMedicine 43, 594–606 (2019).

60. Merrill, C. B. et al. Harnessing changes in open chromatin determined by ATAC-seq to generate insulin-responsive reporter constructs. BMC Genomics 23, 399 (2022).

61. Martinez-Morales, J. R. et al. Differentiation of the vertebrate retina is coordinated by an FGF signaling center. Dev. Cell 8, 565–574 (2005).

62. Peters, M. a & Cepko, C. L. The dorsal-ventral axis of the neural retina is divided into multiple domains of restricted gene expression which exhibit features of lineage compartments. Dev. Biol. 251, 59–73 (2002).

63. Mui, S. H., Kim, J. W., Lemke, G. & Bertuzzi, S. Vax genes ventralize the embryonic eye. Genes Dev. 19, 1249–1259 (2005).

64. Bower, G. & Kvon, E. Z. Genetic factors mediating long-range enhancer–promoter communication in mammalian development. Curr. Opin. Genet. Dev. 90, 102282 (2025).

65. Chepelev, I., Wei, G., Wangsa, D., Tang, Q. & Zhao, K. Characterization of genome-wide enhancer-promoter interactions reveals co-expression of interacting genes and modes of higher order chromatin organization. Cell Res. 22, 490–503 (2012).

66. Kubo, N. et al. Promoter-proximal CTCF binding promotes distal enhancer-dependent gene activation. Nat. Struct. Mol. Biol. 28, 152–161 (2021).

67. Khan, M. A. F. et al. Computational tools and resources for prediction and analysis of gene regulatory regions in the chick genome. genesis 51, 311–324 (2013).

68. Xi, W. & Beer, M. A. Loop competition and extrusion model predicts CTCF interaction specificity. Nat. Commun. 12, 1046 (2021).

69. Fudenberg, G. et al. Formation of Chromosomal Domains by Loop Extrusion. Cell Rep. 15, 2038–2049 (2016).

70. Pachkov, M., Erb, I., Molina, N. & van Nimwegen, E. SwissRegulon: a database of genome-wide annotations of regulatory sites. Nucleic Acids Res. 35, D127–D131 (2007).

71. Castillo, H. et al. Xenopus tropicalis osteoblast-specific open chromatin regions reveal promoters and enhancers involved in human skeletal phenotypes and shed light on early vertebrate evolution. Cells and Development 179, (2024).

72. Hallonet, M., Hollemann, T., Pieler, T. & Gruss, P. Vax1, a novel homeobox-containing gene, directs development of the basal forebrain and visual system. Genes Dev. 13, 3106–3114 (1999).

73. Piccolo, S., Dupont, S. & Cordenonsi, M. The biology of YAP/TAZ: Hippo signaling and beyond. Physiol. Rev. 10.1152/physrev.00005.2014 (2014) doi:10.1152/physrev.00005.2014.

74. Miesfeld, J. B. et al. Yap and Taz regulate retinal pigment epithelial cell fate. Development 142, 3021–3032 (2015).

75. Gestri, G., Bazin-Lopez, N., Scholes, C. & Wilson, S. W. Cell Behaviors during Closure of the Choroid Fissure in the Developing Eye. Front. Cell. Neurosci. 12, 1–12 (2018).

76. Eckert, P., Knickmeyer, M. D. & Heermann, S. In vivo analysis of optic fissure fusion in zebrafish: Pioneer cells, basal lamina, hyaloid vessels, and how fissure fusion is affected by bmp. Int. J. Mol. Sci. 21, (2020).

77. Williamson, K. A. et al. Heterozygous loss-of-function mutations in YAP1 cause both isolated and syndromic optic fissure closure defects. Am. J. Hum. Genet. 94, 295–302 (2014).

78. Holt, R. et al. New variant and expression studies provide further insight into the genotype-phenotype correlation in YAP1-related developmental eye disorders. Sci. Rep. 10.1038/s41598-017-08397-w (2017) doi:10.1038/s41598-017-08397-w.

79. Chow, L., Levine, E. M. & Reh, T. A. The nuclear receptor transcription factor, retinoid-related orphan receptor β, regulates retinal progenitor proliferation. Mech. Dev. 77, 149–164 (1998).

80. Stehlin-Gaon, C. et al. All-trans retinoic acid is a ligand for the orphan nuclear receptor RORβ. Nat. Struct. Mol. Biol. 10, 820–825 (2003).

81. Duan, J., Cai, J., Guo, Y., Bian, X. & Yu, S. ALDH1A3, a metabolic target for cancer diagnosis and therapy. Int. J. Cancer 139, 965–975 (2016).

82. Weldon, S. A. et al. Transcriptomics and chromatin accessibility signatures define the cervical-thoracic boundary along the vertebrate axis. Dev. Biol. 525, 249–258 (2025).

83. Ghinia Tegla, M. G., et al. OTX2 represses sister cell fate choices in the developing retina to promote photoreceptor specification. Elife 9, e54279 (2020).

84. Ceroni, F. et al. Deletion upstream of MAB21L2 highlights the importance of evolutionarily conserved non-coding sequences for eye development. Nat. Commun. 10.1038/s41467-024-53553-2 (2024) doi:10.1038/s41467-024-53553-2.

85. Bhatia, S. et al. Functional Assessment of Disease-Associated Regulatory Variants In Vivo Using a Versatile Dual Colour Transgenesis Strategy in Zebrafish. PLoS Genet. 11, 1–22 (2015).

86. Rieblinger, B. et al. Cas9-expressing chickens and pigs as resources for genome editing in livestock. Proc. Natl. Acad. Sci. U. S. A. 118, (2021).

87. Wong, E. S. et al. Deep conservation of the enhancer regulatory code in animals. Science (1979). 370, (2020).

88. Desmaison, A. et al. Mutations in the LHX2 gene are not a frequent cause of micro/anophthalmia. Mol. Vis. 16, 2847–9 (2010).

89. Moore, J. E. et al. An expanded registry of candidate cis-regulatory elements. Nature https://doi.org/10.1038/s41586-025-09909-9 (2026) doi:10.1038/s41586-025-09909-9.

