## Supplementary figures and images for "The regulatory landscape of optic fissure closure in the vertebrate eye"

### Supplemental Fig S1

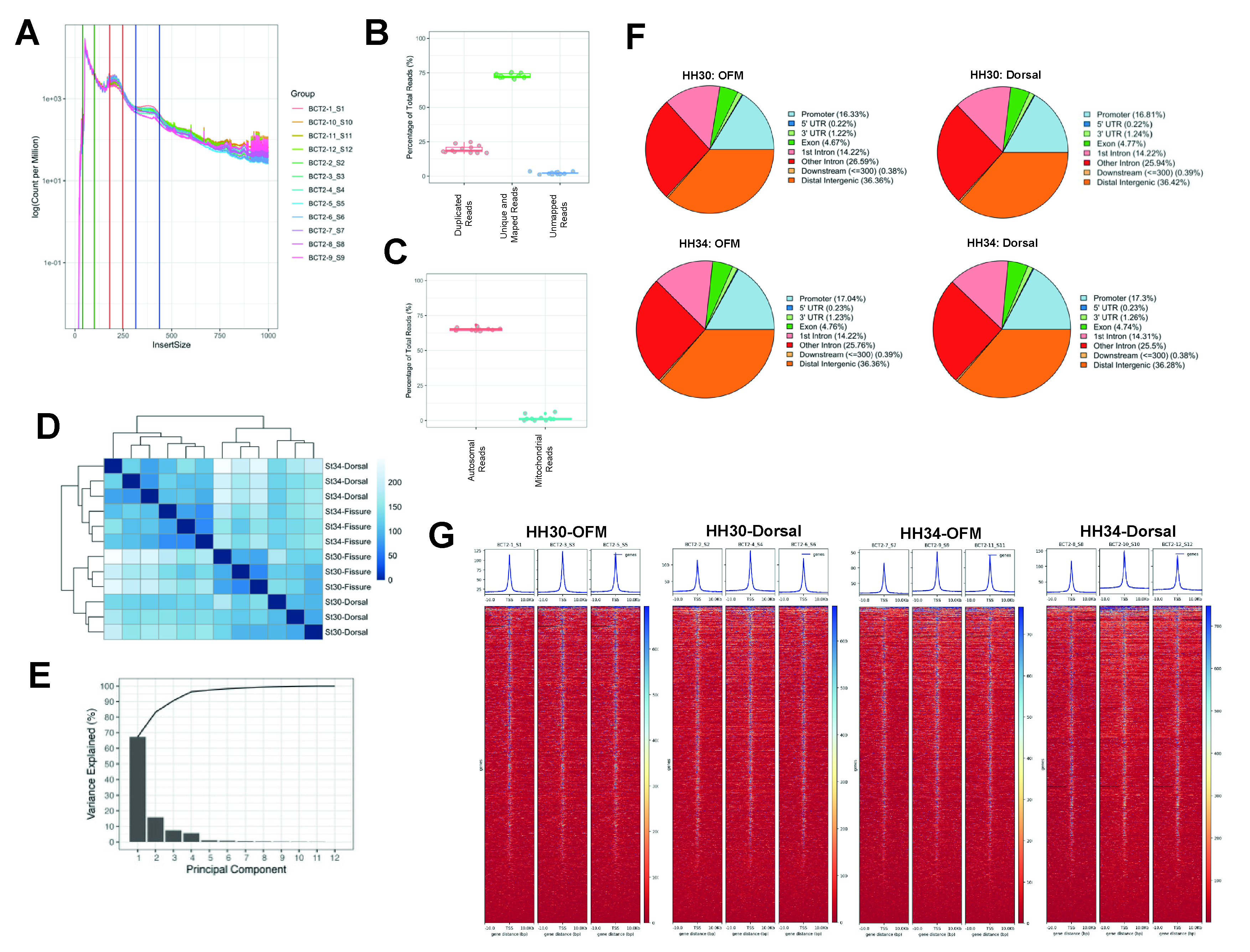

### Supplemental Fig S2

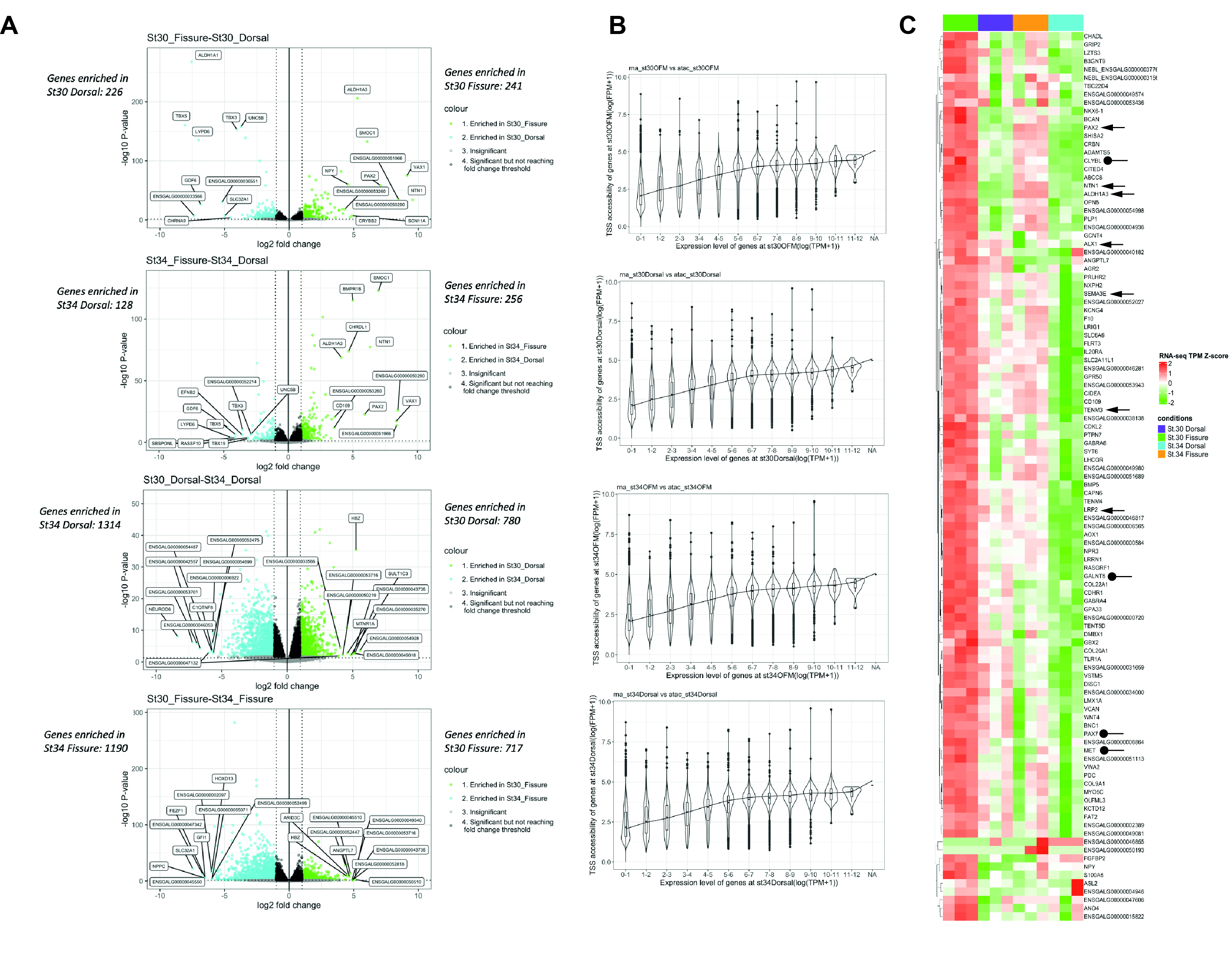

### Supplemental Fig S3

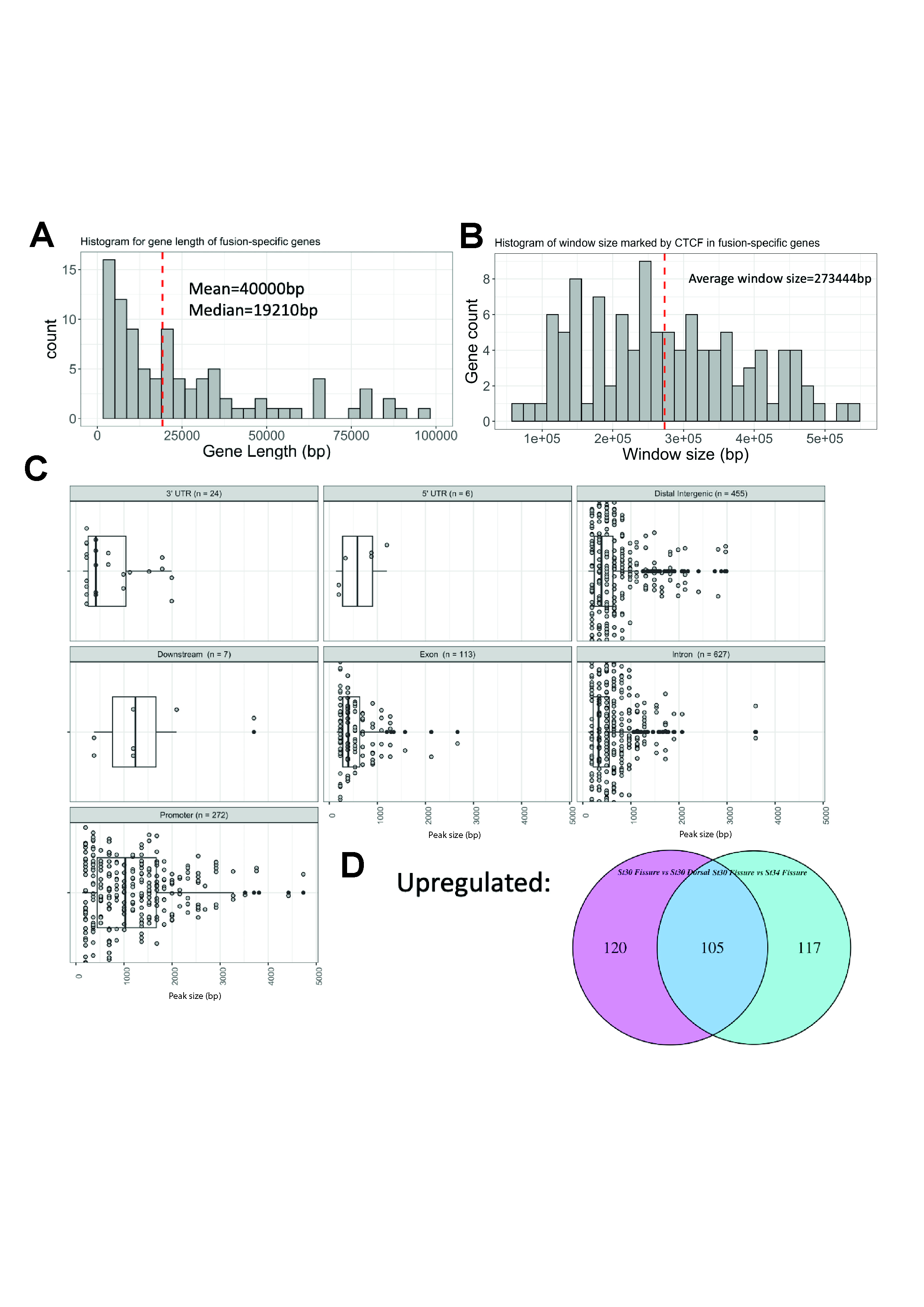

### Supplemental Fig S4

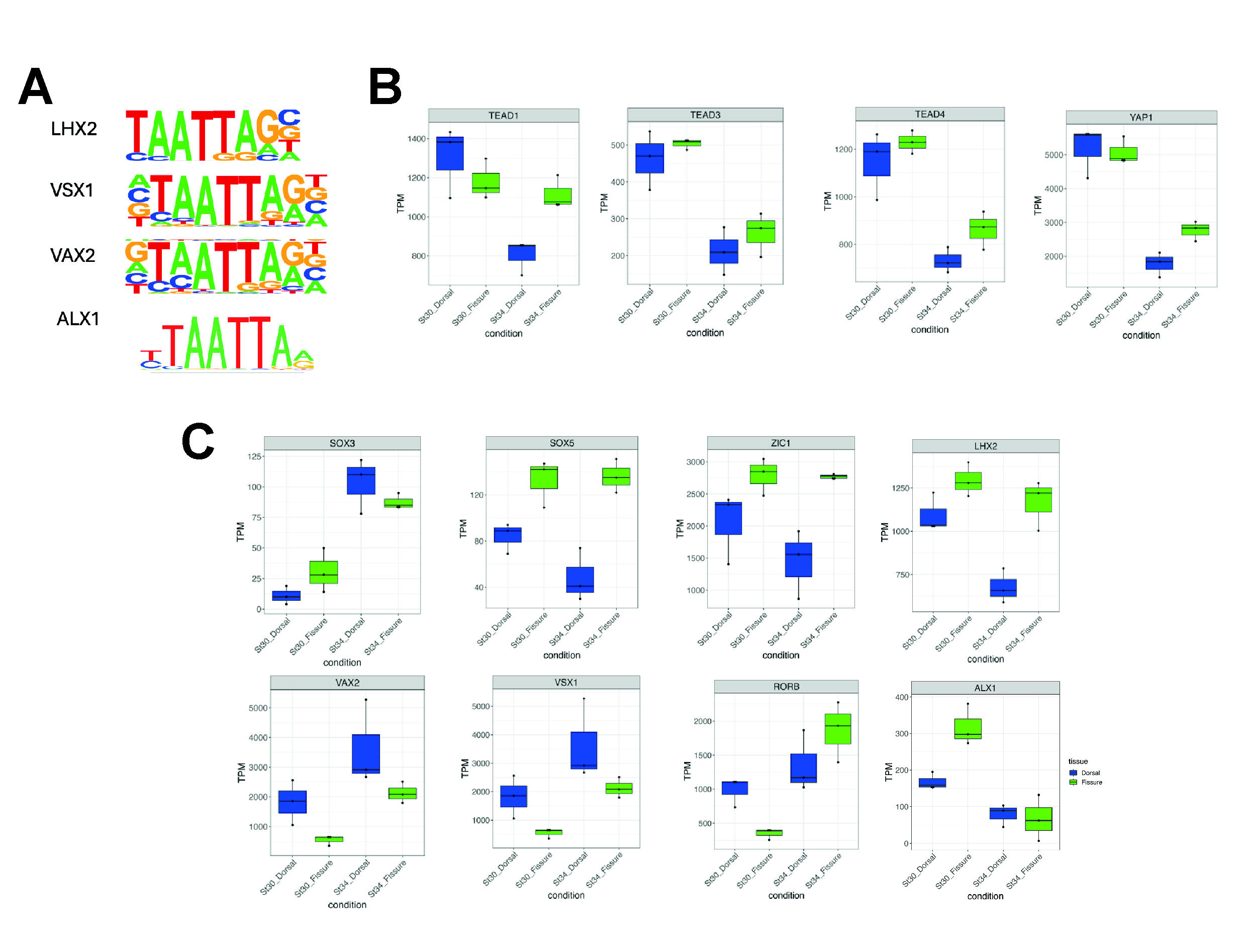

### Supplemental Fig S5

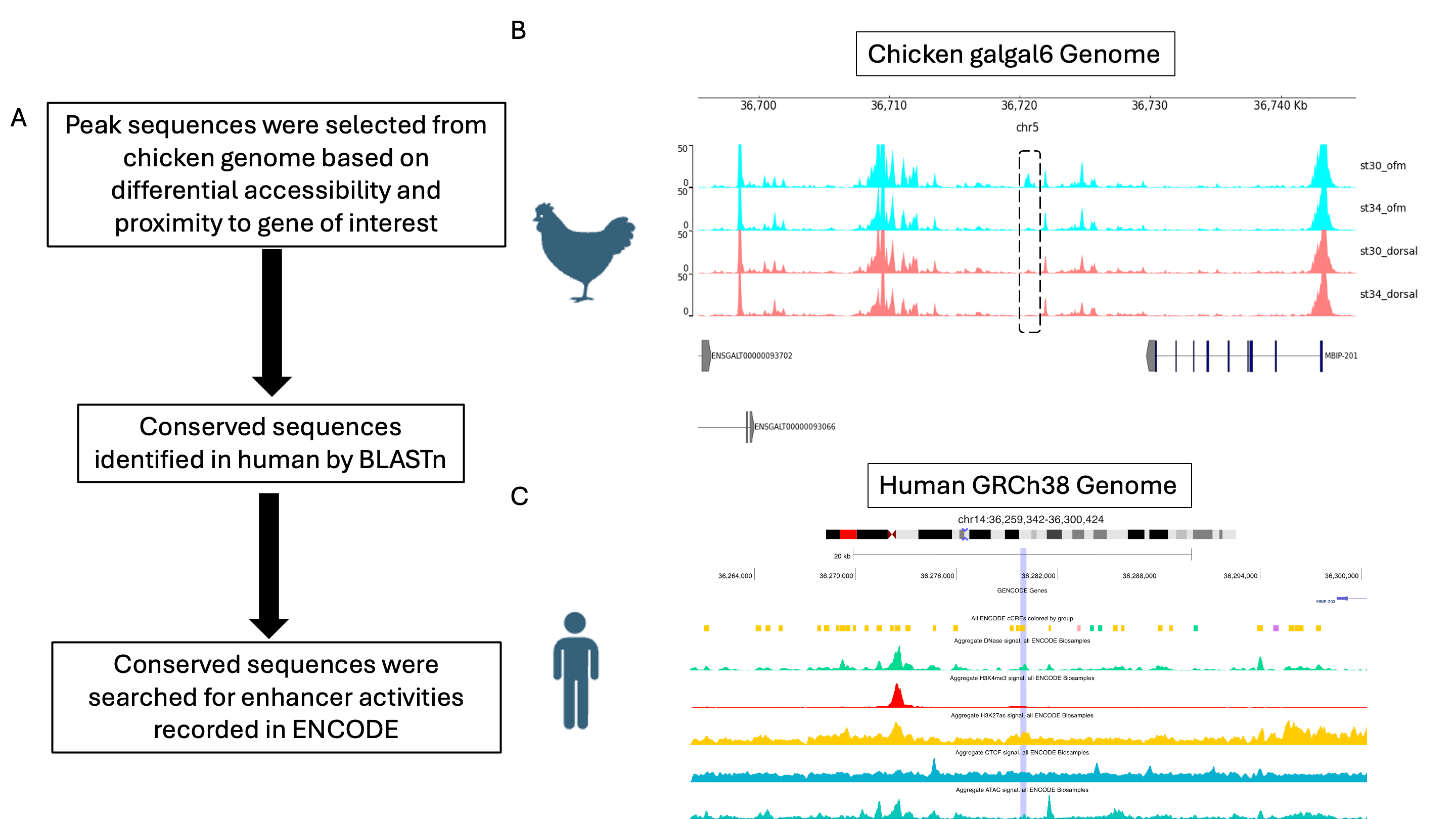
